# Tipping point in *α*-synuclein–membrane interactions: stable protein-covered vesicles or amyloid aggregation

**DOI:** 10.1101/2024.05.01.592005

**Authors:** Katarzyna Makasewicz, Göran Carlström, Olof Stenström, Katja Bernfur, Simon Fridolf, Mikael Akke, Sara Linse, Emma Sparr

**Affiliations:** Division of Physical Chemistry, Department of Chemistry, Lund University, P.O. Box 124, SE-22100 Lund, Sweden; Division of Biophysical Chemistry, Center for Molecular Protein Science, Department of Chemistry, Lund University, SE-22100 Lund, Sweden; Division of Biochemistry and Structural Biology, Center for Molecular Protein Science, Department of Chemistry, Lund University, SE-22100 Lund, Sweden

## Abstract

*α*-synuclein is a neuronal protein implicated in neurotransmitter release. Its function is thought to critically depend on the dynamic equilibrium between free and membrane-bound protein. *α*-synuclein amyloid formation implicated in Parkinson’s Disease was also shown to be modulated by lipid membranes. However, it remains elusive whether *α*-synuclein-related pathology is due to loss-of-function or gain-of-toxic-function. To help address this question, we studied the coupling of the equilibrium between free and membrane-bound *α*-synuclein and membrane-induced amyloid formation – phenomena that are usually treated separately. We present a description of the system on a wide range of length scales and timescales for lipid-to-protein ratio conditions where amyloid formation is either accelerated or inhibited by lipid membranes. We find a clear difference between the dynamics and heterogeneity of the protein-covered membrane interface in the two sets of conditions. In aggregation-accelerating conditions, the membrane interface is dynamic and heterogeneous with rapid exchange between free and membrane-bound protein, and disordered protein segments of varying lengths exposed to solution. All these characteristics of the membrane interface are likely to decrease the free energy barrier for amyloid formation. Conversely, the membrane interface is homogeneous and less dynamic in conditions where amyloid formation is inhibited. Importantly, any factors affecting the equilibrium between free and membrane-bound *α*-synuclein may trigger a change from non-aggregating to aggregating conditions. Altogether, our results highlight a strong coupling of the dynamic equilibrium between the free and membrane-bound *α*-synuclein and membrane-modulated amyloid formation and thus of the physiological function of *α*-synuclein and its aberrant aggregation.

## Introduction

*α*-synuclein is an intrinsically disordered protein enriched in presynaptic termini of neurons. The protein was shown to associate with synaptic vesicles and based on this, it is hypothesized to be involved in regulation of neurotransmitter release.^1^ However, its specific role in the process remains unclear. As a peripheral membrane-binding protein, the function of *α*-synuclein is expected to critically depend on the equilibrium between its free and membrane-bound states. On the other side of the coin, *α*-synuclein is well characterized with respect to its dysfunctional role in neurodegeneration. Its accumulation in the form of amyloid deposits coincides with the loss of dopaminergic neurons in Parkinson’s Disease (PD).^2^ These deposits, referred to as Lewy Bodies, have been shown to contain membrane lipids together with *α*-synuclein amyloid fibrils and other proteins.^3,4^ Moreover, numerous studies have reported that lipid membranes modulate *α*-synuclein amyloid formation.^5–12^ Thus, interactions with lipid membranes play a critical role in both the physiological and pathological function of *α*-synuclein.

Despite extensive studies, it remains unclear whether *α*-synuclein pathology is related to gain-of-toxic function connected to formation of amyloid fibrils, loss of function connected to depletion of nonfibrillar protein, or to a combination of the two effects.^13^ Since interactions with lipid membranes are key to both processes, detailed characterization of the coupling of the equilibrium between free and membrane-bound protein and amyloid formation in presence of membranes may help revealing the basic mechanisms governing the physiological *α*-synuclein–lipid co-assembly, and how this can be turned into pathological protein–lipid amyloid co-aggregates. Previous studies already indicate the presence of such coupling: lipid membranes were shown to either accelerate or inhibit *α*-synuclein amyloid formation depending on the relative amounts of proteins and lipids, with acceleration occurring under conditions of excess protein, where initially free and membrane-bound protein coexist, and inhibition occurring under conditions of excess membrane, where virtually all protein is associated with the membrane.^5^ Moreover, all known PD-related mutations are located in the membrane-binding region of the *α*-synuclein sequence,^1^ suggesting that the acceleration of aggregation by membranes may be due to the perturbation of protein-membrane interactions and not necessarily to the higher aggregation propensity of the disease mutants. Thus, we take a viewpoint of having one system that can develop in different directions (stable protein-coated vesicles or amyloid formation) rather than treating the amyloid formation and the adsorption of protein to the membrane as separate processes, which is commonly done in the literature. The consideration of the coupling between the molecular events in these two directions and the conditions that direct the co-assembly is expected to shed light on the molecular origins of *α*-synuclein-related pathology.

In this work, we provide a detailed description of *α*-synuclein interactions with membranes across a wide range of timescales and length scales for conditions where the protein is present in excess as well as conditions where there is an excess of lipid membranes (Figure 1), expressed in terms of the relative amounts of lipids and proteins; the lipid-to-protein ratio (L/P). First, we determine the conditions where the presence of lipid membranes leads to either acceleration or inhibition of *α*-synuclein amyloid formation using thioflavin T (ThT) aggregation kinetics assay. Next, we characterize the system in terms of relative amounts of free and bound protein across the studied L/P range employing native agarose gel electrophoresis, circular dichroism spectroscopy, and Pulsed-Field Gradient (PFG) diffusion NMR spectroscopy. Finally, we characterize in detail the membrane-bound state and the exchange dynamics between free protein and different membrane-bound states using ^1^H-^15^N Heteronuclear Single Quantum Coherence (HSQC) and PFG diffusion NMR. Thus, we study the system on three levels: the macroscopic, the colloidal, and the molecular level (Figure 1).

**Figure 1.**
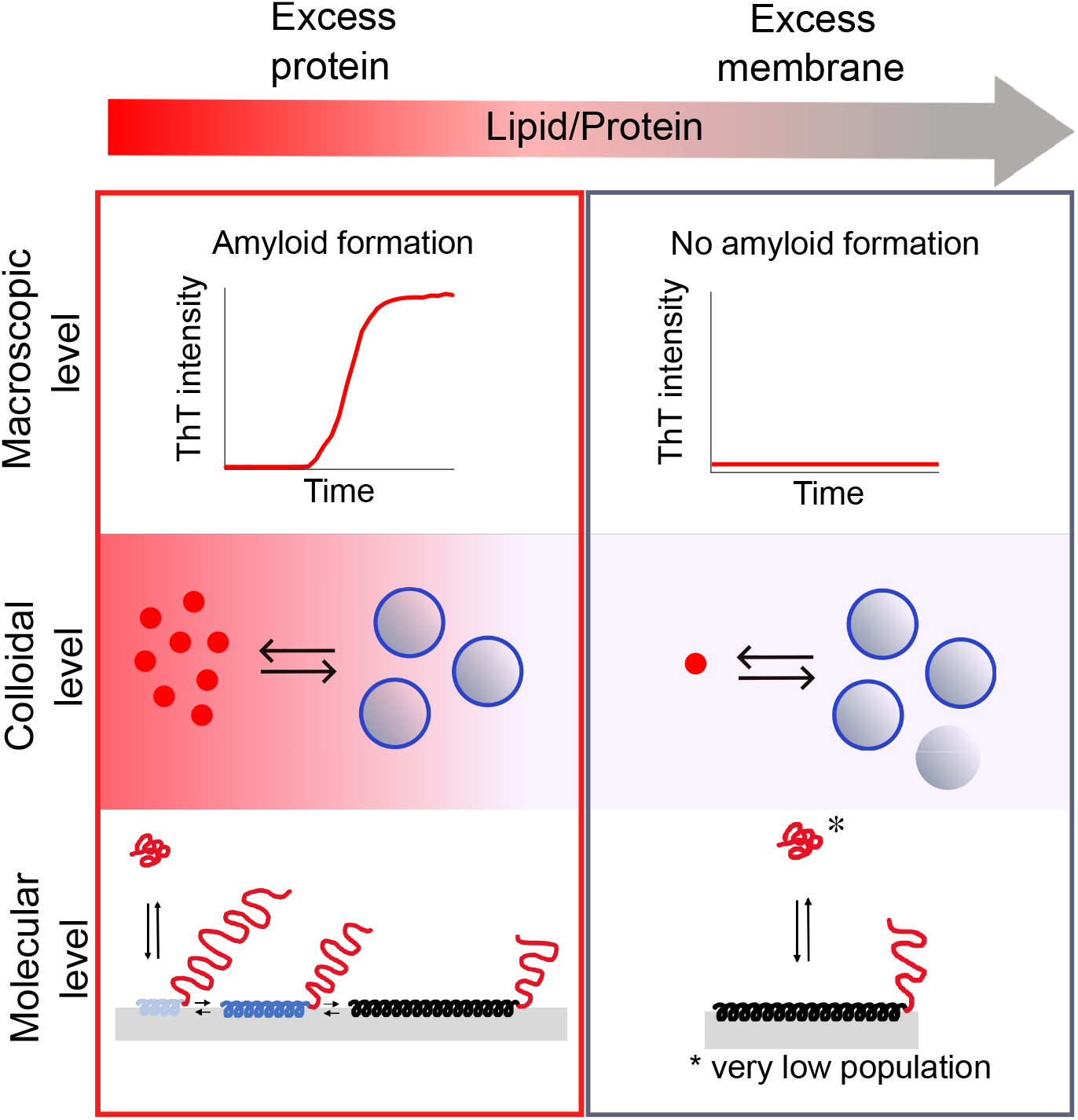
*α*-synuclein interactions with lipid membranes on macroscopic, colloidal and molecular levels as a function of the relative amounts of lipids and proteins, quantified by the lipid-to-protein (L/P) ratio. The macroscopic level refers to the presence or absence of amyloid formation. The colloidal level corresponds to the relative amounts of free and membrane-bound protein, while the molecular level corresponds to the different membrane-bound states present in equilibrium with free protein in solution and to the timescale of the exchange between the different states. Red and blue colors indicate free and membrane-bound protein, respectively. The background of the colloidal level and the L/P ratio axis are colored according to the amount of free protein.

Our results reveal clear differences in the behavior of the system on a molecular level between conditions where *α*-synuclein amyloid formation is accelerated (low L/P ratio) or inhibited (high L/P ratio) by lipid membranes. Under conditions where amyloid aggregation is accelerated, the protein-covered membrane interface is highly dynamic and heterogeneous, while under aggregation-inhibiting conditions, the membrane interface is far less dynamic with a more homogeneous structure. Rapid exchange of free monomer with the membrane-bound state and the possibility of monomer interaction with the exposed aggregation-prone segments of membrane-associated proteins may contribute to lowering of the free energy barrier for amyloid formation at low L/P ratio conditions. On the other hand, slow exchange dynamics and the presence of a relatively stable fully membrane-bound *α*-synuclein state at high L/P ratio may give rise to the either thermodynamic or kinetic stability of the system at these conditions.

Under aggregating conditions, partly membrane-bound states are present, which interact with the membrane less strongly as compared to the fully membrane-bound state. Any factors that affect protein-membrane interaction strength, such as solution conditions, membrane properties, protein mutations or post-translational modifications (PTMs) are likely to shift the system from aggregating to non-aggregating conditions, or vice versa, at a fixed L/P ratio. Thus, our results highlight a strong coupling of the dynamic equilibrium between the free and membrane-bound *α*-synuclein and membrane-modulated amyloid formation, which is highly relevant for the design of therapeutic approaches for *α*-synuclein-related neurode-generative diseases.

## Results

### *α*-synuclein and lipid vesicles on a macroscopic level

First, we determined the effect of DOPC:DOPS 7:3 small unilamellar vesicles (SUVs) on *α*-synuclein amyloid formation in a broad L/P range (10-250). The aggregation was monitored via the thioflavin T (ThT) fluorescence intensity for samples containing only *α*-synuclein or both *α*-synuclein and negatively charged SUVs. 20 *µ*M *α*-synuclein with 20 *µ*M ThT in 10 mM MES pH 5.5 was incubated with increasing content of SUVs in PEGylated polystyrene plates under quiescent conditions at 37^°^C for 18 days. Figure 2A shows the aggregation half-times plotted as a function of the L/P ratio. For samples containing only *α*-synuclein and no vesicles, we detected aggregation in 4 out of 10 replicates with the halftimes ranging from 14 to 16.5 days. More rapid amyloid formation was observed in the presence of vesicles in the L/P range 10-90. Importantly, for the samples at L/P 10-40, we detected aggregation in all 10 replicates, while no aggregation in the time-frame of the experiment was detected for samples at L/P ≥ 100. The maximum ThT intensity observed in our assay does not depend on the total lipid concentration in the sample (Figure S1), which is consistent with previous reports that the presence of lipids does not affect the ThT fluorescence intensity. ^14,15^

**Figure 2.**
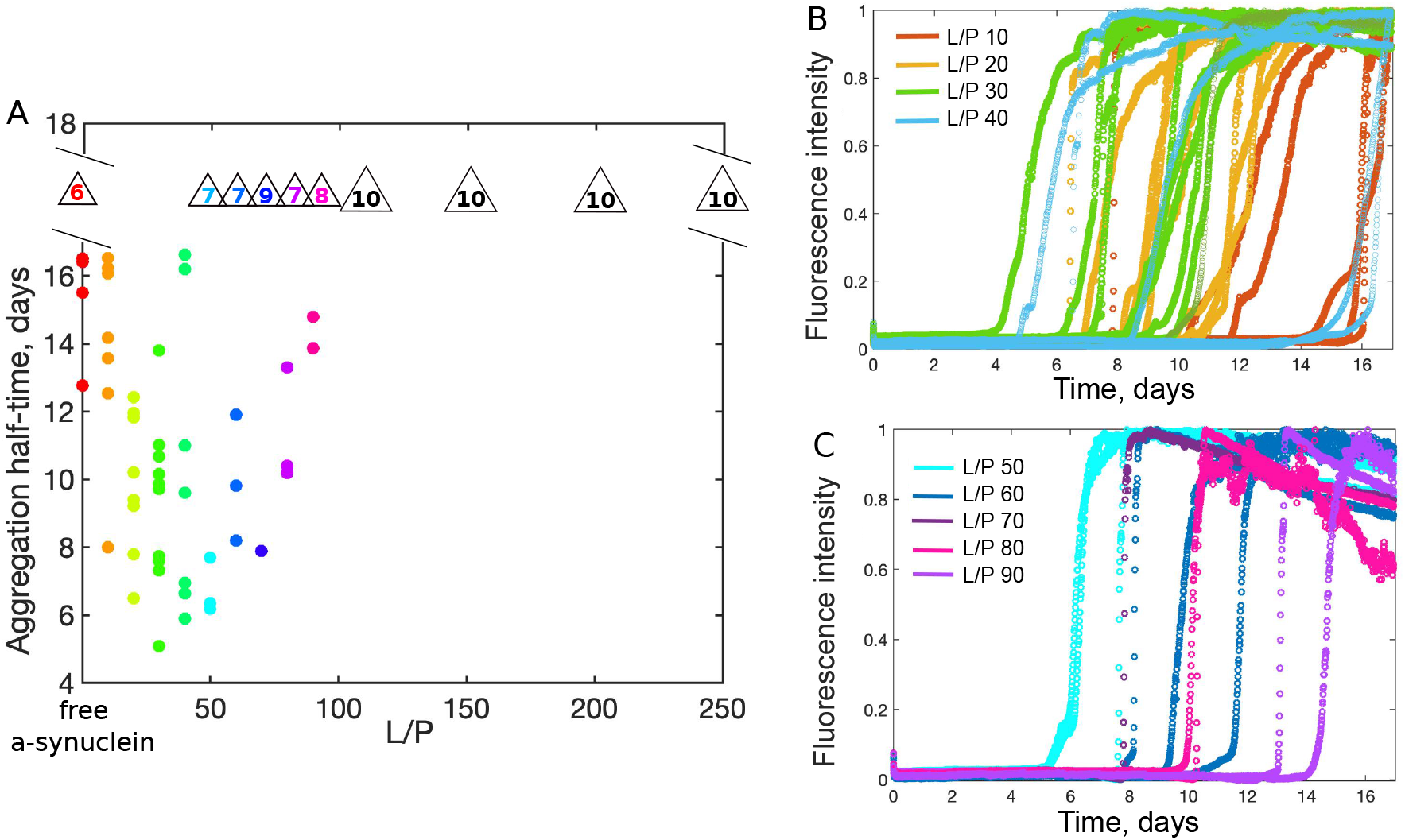
Aggregation kinetics of 20 *µ*M *α*-synuclein in 10 mM MES, pH 5.5 in the presence of DOPC:DOPS 7:3 SUVs in the L/P ratio range 10-200 at 37 ^°^C studied using thioflavin T (ThT) fluorescence. A) The aggregation half-times (the time at which the ThT fluorescence signal reaches half of the maximum intensity) in days as a function of L/P ratio. The number of replicates for which no aggregation was detected is indicated in the triangles above the data points. The time frame of the experiment was 18 days and only repeats for which aggregation was observed within this time-frame are shown. B,C) Normalized ThT intensity versus time for samples at L/P ratios 10-40 (B) and 50-90 (C). No ThT fluorescence above the baseline was detected for samples at L/P ≥ 100.

### *α*-synuclein and lipid membranes on a colloidal level

#### Free and membrane-bound α-synuclein based on electrophoretic mobility and circular dichroism

We first characterized the equilibrium between free and vesicle-bound *α*-synuclein by monitoring the electrophoretic mobility of the species present in the sample. An external electric field was applied to samples deposited in the wells of an agarose gel to monitor the migration of the sample components towards the oppositely charged electrode with the speed governed by their size and charge. We used *α*-synuclein labelled with AlexaFluor-488 and vesicles containing a small fraction of a fluorescent lipid analogue (DHPE-Atto-647). The protein concentration was held constant at 25 *µ*M, while the lipid concentration was varied to cover the L/P ratios from 0 to 250. The buffer conditions used were 50 mM MES pH 5.5 to ensure sufficiently high ionic strength and thereby current.

Free protein, free vesicles as well as vesicles with associated protein were observed to migrate in the gel towards the cathode, indicating that the overall charge of these species is negative. In the samples at L/P 10-50, a strong band corresponding to the free protein is observed (Figure 3A). The band is weaker at L/P 100 and not discernible at L/P ≥ 150. This implies that virtually all protein is associated with vesicles at L/P 150 and higher. We attribute the second less intense band present in the sample containing only free protein and in the L/P 10-50 samples to charge inhomogeneity of the covalently-attached fluorescent dye. An orange diffuse band with higher electrophoretic mobility than free protein corresponds to vesicles with associated protein. Thus, protein-decorated vesicles migrate through the gel at higher speed than the free protein, which can be ascribed to a more negative overall charge. The high fluorescence intensity of the free protein band in the samples at low L/P precludes more quantitative analysis of the amount of free protein. The red fluorescence from the vesicles is indiscernible at the lowest L/P ratios due to low concentration of lipids in these samples. Upon binding to the lipid membrane, the segment of *α*-synuclein spanning residues 1-100 can undergo a conformational transition from random coil to *α*-helix.^16–18^ Therefore, methods sensitive to conformational changes, such as circular dichroism (CD) spectroscopy, have been employed as an indirect tool to study *α*-synuclein binding to lipid membranes. Figure 3B shows CD spectra of free *α*-synuclein and *α*-synuclein in presence of SUVs for L/P ratios ranging between 10-250. When SUVs are titrated into the solution of *α*-synuclein, the mean residue ellipticity (MRE) at 222 nm becomes more negative (Figure 3C), which is indicative of an increase in the *α*-helical content. MRE at 222 nm changes linearly with increasing lipid concentration in the L/P range 10-160 and reaches a stable value above L/P 160. This implies that the average number of residues in *α*-helical conformation increases linearly with L/P ratio up to L/P 160. Above L/P 160, the maximum *α*-helical content is attained, implying that all protein molecules are associated with the vesicles, in line with the agarose gel electrophoresis results. We emphasize that the CD signal is a time and bulk average and the decrease in the CD signal at 222 nm (i.e. more negative signal) may be due to a larger number of protein molecules containing a short *α*-helical segment, or smaller number of protein molecules containing a long *α*-helical segment, as well as any intermediate case. The analysis of the CD data does not permit differentiation between these scenarios.

**Figure 3.**
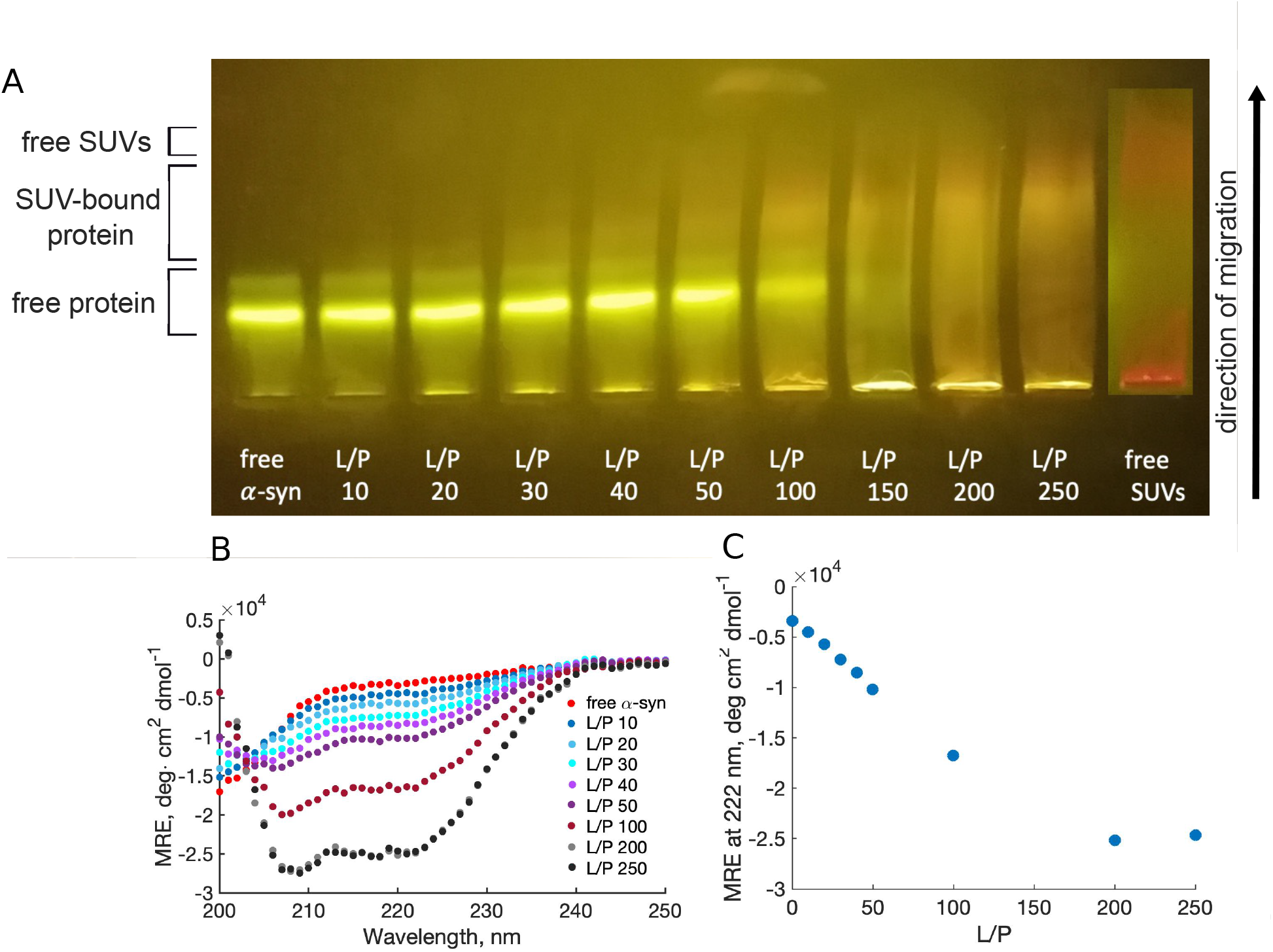
Estimation of the amount of free and membrane-bound *α*-synuclein across a broad L/P ratio range. A) Agarose gel electrophoresis of samples containing *α*-synuclein-N122C labelled with Alexa Fluor 488 Maleimide (*α*-synuclein-488) and DOPC:DOPS 7:3 SUVs containing 2% DHPE-647. In all samples, the *α*-synuclein-488 concentration was 20 *µ*M, while the concentration of lipids was varied to reach a given L/P ratio. The buffer conditions were 50 mM MES pH 5.5. The protein appears yellow, while the vesicles appear red. The lipid concentration is too low for the red fluorescence to be discernible at low L/P ratio. Electrophoresis is run in horizontal mode, and the samples migrate from the anode (bottom of the gel image) towards the cathode (top of the gel image). B,C) Conformational change of *α*-synuclein upon association with lipid membranes studied by means of circular dichroism (CD) spectroscopy. B) CD spectra of *α*-synuclein alone and in presence of DOPC:DOPS 7:3 SUVs in the L/P ratio range 10-250. C) Mean residue ellipticity at 222 nm as a function of L/P ratio extracted from the CD spectra shown in B.

#### Dynamic equilibrium between free and membrane-associated *α*-synuclein studied with PFG diffusion NMR spectroscopy

PFG NMR spectroscopy allows one to study the dynamic equilibrium between free and membrane-associated protein and extract the timescale of exchange between the two states and their relative populations. If the studied molecule can exist in two states characterized by different diffusion coefficients, the possibility of detecting two diffusing components in the PFG experiment will depend on the timescale of the exchange between the two states relative to length of the diffusion period (diffusion time) employed in the experiment.^19^ If the exchange occurs on a timescale that is much shorter than the diffusion time, then the observed signal decay will be monoexponential with the decay constant being the population-averaged diffusion coefficient. If the exchange occurs on a timescale that is longer than the diffusion time, then the observed signal decay will be a sum of exponentials, each with a characteristic diffusion coefficient.

PFG diffusion NMR experiments were performed on samples containing 200 *µ*M *α*-synuclein with or without SUVs at L/P 50, 100 and 200 in 10 mM MES buffer, pD 5.5 prepared in D_2_O. Prior to PFG experiments, we identified the resonances in a 1D ^1^H spectrum originating from the protein, which do not overlap with the resonances from the lipids. As shown in the ^1^H spectra in Figure 4A, the only *α*-synuclein peaks that do not overlap with the lipid peaks are those at 6.9 and 7.2 ppm, which originate from the aromatic side chains.

**Figure 4.**
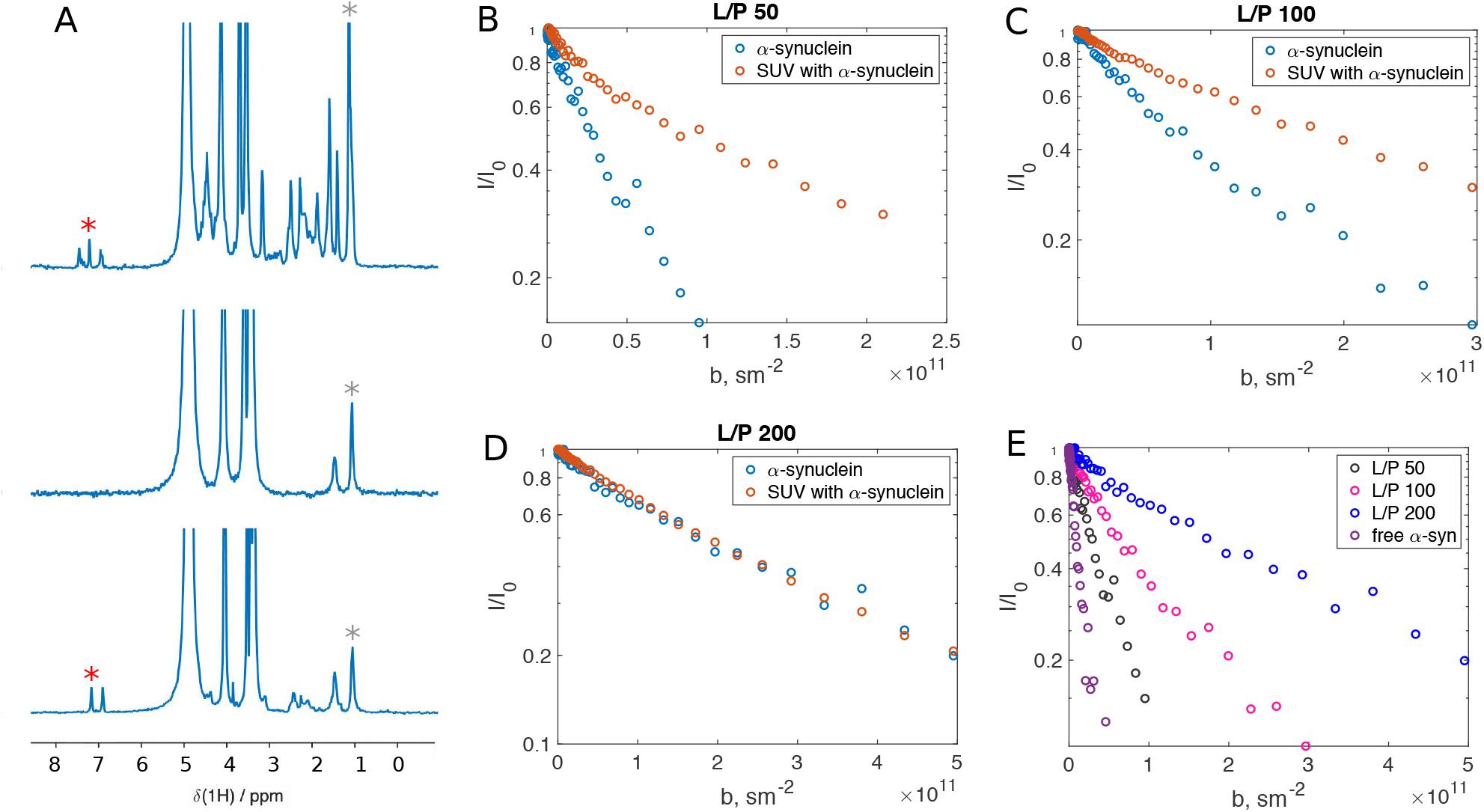
Quantitative analysis of the equilibrium between free and membrane-bound *α*-synuclein at different L/P ratio conditions with PFG Diffusion NMR spectroscopy. A) ^1^H NMR spectra of free *α*-synuclein (top), protein-free small unilamellar vesicles composed of DOPC:DOPS 7:3 (middle) and *α*-synuclein with SUVs at L/P 100 (bottom). The decay of the intensities of the peak at 7.2 ppm (from *α*-synuclein, red *) and 1 ppm (from lipids and *α*-synuclein or protein-free lipids, grey *) was monitored to study the translational diffusion in the PFG diffusion NMR experiment. B, C, D) Intensity decay curves of *α*-synuclein (7.2 ppm peak, blue circles) and SUVs+*α*-synuclein (1 ppm peak, orange circles) at L/P 50, 100 and 200. The peak intensities at increasing gradient strength were normalized with respect to the peak intensity in the absence of the gradient field and plotted as a function of *b*=(*γGδ*)^2^(Δ − *δ/*3), where *γ* is the proton gyromagnetic ratio (rad/s/T), *G* is the gradient strength (T/m), *δ* is the gradient pulse length and Δ is the total diffusion time (s). E) The intensity decay curves for the peak corresponding to *α*-synuclein (7.2 ppm) for samples containing free *α*-synuclein or *α*-synuclein and SUVs at L/P 50, 100 and 200. For all samples the diffusion time in the pulse sequence was kept constant at 50 ms. The maximum gradient strength was optimized for each sample to ensure sufficient decay of the peaks corresponding to *α*-synuclein.

Figures 4B-D show the intensity decay of the *α*-synuclein peak (7.2 ppm) and the lipid+*α*-synuclein peak (1 ppm) in the samples at L/P 50, 100 and 200, respectively. In all cases, the signal decay follows a single exponential. The fact that at L/P 200 the decay curves of the protein and lipid+protein signals overlap corroborates the CD and agarose gel electrophoresis results showing that essentially all protein molecules are associated with lipid vesicles under these conditions. The intensity decay curves for the *α*-synuclein peak in all samples are plotted in Figure 4E. The decrease in the decay rate with increasing L/P ratio reflects an increase in the fraction of bound protein.

The monoexponential decays of the *α*-synuclein peaks (7.2 ppm) in the samples at L/P 50 and 100, imply that the lifetimes of the free and membrane-bound states are much shorter than the diffusion time in the pulse sequence (50 ms). ^19^ In other words, during the time frame of 50 ms, a large number of association/dissociation events takes place. Accordingly, at these L/P ratio conditions, the exchange rate between the free and membrane-bound protein, 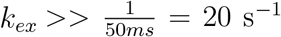. Since the intensity decay curves follow a single exponential decay, the extracted diffusion coefficients are population averages of the free and vesicle-associated protein.^19^ Thus, they can be used to estimate the fraction of vesicle-bound *α*-synuclein at each L/P ratio. The calculated fractions of bound protein are 74%, 95% and 100% at L/P 50, 100 and 200, respectively (for details see Supplementary Information and Table S1).

### *α*-synuclein and lipid membranes on a molecular level

#### Characterization of the membrane-bound states

It has been reported that *α*-synuclein can engage in membrane interaction with sequence stretches of different length.^20,21^ Thus, in order to characterize the membrane-bound state of *α*-synuclein, an experimental technique reporting on which segments of the protein are associated with the membrane at given conditions is necessary. To this end, we carried out ^1^H-^15^N HSQC experiments on samples containing *α*-synuclein isotope-labelled with ^15^N and SUVs at different L/P ratios (Figure 5 and Figure S2). The ^1^H-^15^N HSQC spectrum of free *α*-synuclein contains cross-peaks corresponding to most of the residues in the protein’s sequence apart from prolines, which lack an amide proton (Figure 5A, blue). Upon addition of SUVs (Figure 5A, red) some of the peaks disappear and many appear with reduced intensity. Figure 5B shows the peak intensities from the ^1^H-^15^N HSQC spectra at 20^°^C for L/P 50, 100 and 200, normalized to the intensity of the corresponding peaks in the spectrum of *α*-synuclein without any SUV present. When the peak intensity is analyzed as a function of residue number in the protein sequence, it becomes apparent that the invisible peaks belong to the residues in the membrane-binding segment (residues 1-100).

**Figure 5.**
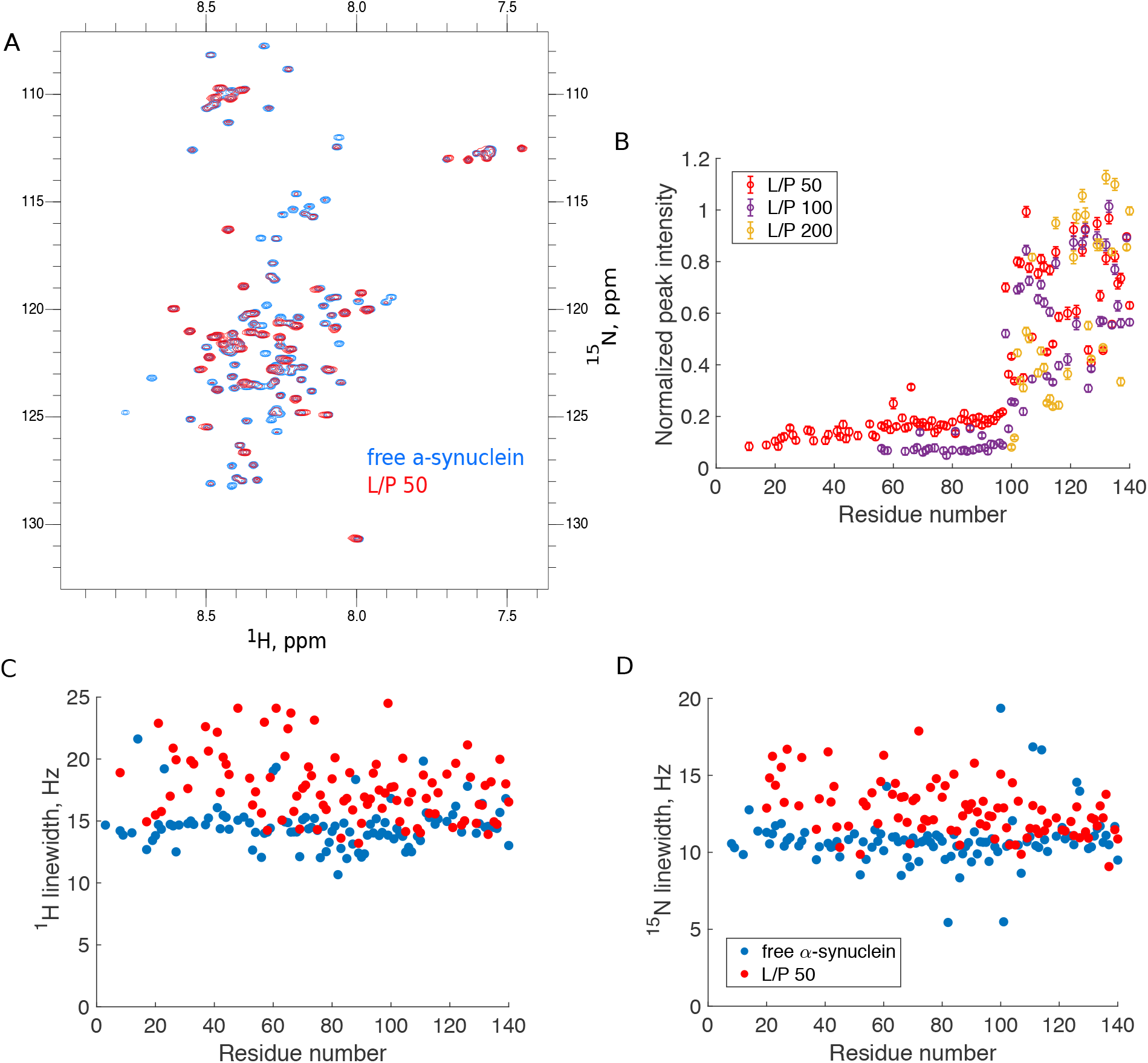
*α*-synuclein association with DOPC:DOPS 7:3 SUVs studied with ^1^H-^15^N HSQC NMR. A) The spectrum of free *α*-synuclein (blue) overlaid with the spectrum of *α*-synuclein with SUVs at L/P 50 (red) at 20^°^C. B) Peak intensities in the samples containing *α*-synuclein and SUVs at L/P 50, 100 and 200 at 20^°^C normalized with respect to the intensity of the corresponding peaks in the free *α*-synuclein spectrum and plotted as a function of the residue number in the *α*-synuclein sequence. The resonances for residues 1-7 in the presence of SUVs could not be reliably analyzed due to peak overlap. C,D) Linewidths of the ^1^H (C) and ^15^N (D) resonances in the HSQC spectrum of *α*-synuclein with SUVs at L/P 50 (red) and in the absence of SUVs (blue) at 20^°^C. Peak volumes and linewidths were evaluated using the program suite PINT.^22,23^

Due to slow rotational diffusion of the vesicle, residues bound to the membrane experience very high transverse relaxation rates, *R*_2_, which results in broadening of the peaks beyond detection. Sufficiently rapid exchange between free and vesicle-bound protein may also lead to broadening of the resonances beyond detection. Residues that are not membrane-bound but extend away from the vesicle surface experience much shorter correlation times than the vesicle itself due to the inherent flexibility of the intrinsically disordered protein chain. Thus, the observable but attenuated signals reflect the fraction of protein that is not adsorbed to the membrane with a given sequence segment, including both the completely free protein and the extended part of the vesicle-bound protein.

The signals from residues 101-140 remain clearly observable at all analyzed L/P ratios, albeit with varying attenuation, implying that this segment does not associate to any greater extent with the lipid membrane. Five proline residues in this segment likely affect the peak intensities of surrounding residues due to increased chain stiffness and cis-trans isomerization of the peptidyl-prolyl peptide bond, which might also affect transient association with lipids.^20^

Comparison of the trends of normalized peak intensity (Figure 5B) as a function of residue number provides insights into what happens at the membrane surface at the different L/P ratio conditions. At L/P 50, residues in the N-terminal segment are either invisible (D2, V3, L8, S9 and K12) or not possible to analyze due to peak overlap, with A11 showing very weak intensity. The visible but attenuated resonances belong to residues 17-100, which have peak intensities of approximately 15% compared to the protein sample without SUVs. At the face of it, this observation might imply that 85% of the protein molecules are bound to the vesicle with the full segment engaged in membrane binding (i.e., residues 1-100 are in contact with the membrane). However, this conclusion does not take into account the effect of line-broadening due to exchange between the free and vesicle-bound species; we return to this issue below. The observable resonances have chemical shifts virtually identical to those of the vesicle-free sample and display linewidths broadened by a few Hz (Figure 5C).

In the HSQC spectrum of *α*-synuclein at L/P 100, the stretch of residues not clearly observable above the noise level extends from 1 to 55 (Figure 5B). In the segment 56-100, the peak heights are approximately 7% compared to the free protein. At L/P 200, on the other hand, no resonances in the membrane-binding region are observable, which implies that virtually all protein molecules are membrane-associated throughout the segment 1-100 and the concentration of the free protein is too low to be detected.

Altogether, the data in Figure 5B imply that at L/P ratio of 50, where protein is present in excess (Figure 3A) *α*-synuclein engages in membrane interactions with different binding modes where either a short (residues 1-16) or a long (residues 1-100) segment is associated with the membrane. At L/P 100, again two different binding modes are detected, with either an intermediate (residue 1-55) or a long (1-100 residues) sequence segment is associated with the membrane. At the highest L/P of 200, where all protein is bound (Figure 3A), the only present binding mode spans the full membrane-binding segment of the protein (residues 1-100).

We also acquired ^1^H-^15^N HSQC spectra at L/P 10, 20, 40 and 50 at 37^°^C (Figure S3). The attenuation of the peaks follows the same pattern as observed at 20^°^C, but the changes occur in a different L/P ratio range (Figure S4). The attenuation of the signal at L/P 50 at 20^°^C is identical to the attenuation at L/P 10 at 37^°^C. The same holds for L/P 100 (20^°^C) versus L/P 20 (37^°^C), and for L/P 200 (20^°^C) versus L/P 40 (37^°^C). These observations are consistent with the temperature dependence of *α*-synuclein binding to lipid membranes determined from CD spectroscopy (Figure S5). An increase in the protein-membrane interaction strength with increasing temperature is consistent with hydrophobic interaction of the amphipathic *α*-helix with the hydrophobic core of the membrane playing a crucial role in binding.

#### Characterization of the exchange dynamics between free and membrane-bound *α*-synuclein

The results of the HSQC and diffusion NMR experiments permit us to estimate the timescale of the exchange process between free and membrane-bound *α*-synuclein. From the diffusion NMR experiment, we conclude that the exchange rate constant between these two states is *k*_1*ex*_ *>* 20 *s*^−1^. Importantly, since all *α*-synuclein membrane-bound states—regardless of the length of the associated protein segment—are characterized by the diffusion coefficient of the vesicle, the exchange process probed in the diffusion NMR experiment is between the free protein and any of the possible membrane-bound states.

The effect of exchange processes on the HSQC spectrum depends on whether the exchange falls in the slow, intermediate or fast regime with respect to the NMR chemical shift timescale,^24^ and on the difference in the transverse relaxation rates, *R*_2_, of the two states. In the simplest case, when the two exchanging states are characterized by similar *R*_2_, the conditions for the slow, fast and intermediate exchange regime are *k*_1*ex*_ *<<* Δ*ω, k*_1*ex*_ ≈ Δ*ω* and *k*_1*ex*_ *>>* Δ*ω*, respectively, where Δ*ω* is the difference in chemical shifts between the two states.^24^ Thus, for a given Δ*ω*, a low *k*_1*ex*_ gives rise to an NMR spectrum with two resonances at their characteristic chemical shifts (slow exchange regime). Increasing *k*_1*ex*_ causes broadening and merging of the two resonances (intermediate exchange regime), and further increasing *k*_1*ex*_ leads to reappearance of a sharp line at a chemical shift determined by the relative populations of the two states and their intrinsic chemical shifts (fast exchange regime) (Figure S6A). Based on this type of analysis, the fact that the observable resonances from the membrane-bound segment of *α*-synuclein in the presence of vesicles display unchanged chemical shifts with respect to chemical shifts in the spectrum of vesicle-free protein, Bodner et al. concluded that the exchange process between free and membrane-bound *α*-synuclein is slow on the NMR chemical shift timescale.^20^ However, when the difference in the transverse relaxation rates of the two exchanging states (Δ*R*_2_) is large, the exchange regime is determined by the value of *k*_1*ex*_ relative to |Δ*ω* + *i*. Δ*R*_2_|.^25^ We estimate the *R*_2_ values for the free and membrane-bound protein to be approximately 10 *s*^−1^ and 39,000 *s*^−1^ (details in SI). NMR spectra for spins exchanging between two sites with such *R*_2_ values are plotted in Figure S6B. In our case, the large Δ*R*_2_ will dominate the expression |Δ*ω* + *i*. Δ*R*_2_|. The effect of Δ*R*_2_ on the appearance of the NMR spectrum with two exchanging species is plotted in Figure S7: regardless of the value of *k*_1*ex*_, the resonance of free protein exchanging with vesicle-bound protein will have unchanged chemical shift compared to the free protein in the absence of SUVs. Thus, due to a large Δ*R*_2_ between the free and vesicle-bound states of *α*-synuclein, the exchange regime cannot be estimated based on the NMR chemical shift of the observable residues.

In order to extract exchange rate constants for the present system we need to take a different approach. The fact that residues 1-16 are unobservable in the HSQC spectrum at L/P 50 allows us to extract boundaries on *k*_1*ex*_ for the exchange process between free and the 1-16 membrane-bound state. The diffusion NMR experiments (Figure 4) show that 26% of the protein is free in solution at these conditions. We can thus conclude that residues 1-16 of the free protein fraction are unobservable in the HSQC spectra because of exchange-broadening beyond detection. To estimate the value of *k*_1*ex*_ between free protein and 1-16 membrane-bound state, we calculated NMR spectra for a species exchanging between two states with *R*_2_ of 10 *s*^−1^ (free protein) and 39,000 *s*^−1^ (membrane-bound protein). We find that the resonances become exchange-broadened beyond detection when *k*_1*ex*_ ≥ 300 *s*^−1^ (Figure S8A), in agreement with the results of the diffusion experiments, which yielded *k*_1*ex*_ *>* 20 *s*^−1^. The same explanation applies to the segment 1-55 at L/P 100: free protein exchanges with the 1-55 membrane-bound state with *k*_1*ex*_ ≥ 300 *s*^−1^.

The extent of line broadening of the remaining observable resonances from the membrane-binding region (residues 17-100 at L/P 50 or residues 56-100 at L/P 100) permits us to estimate the timescale of the exchange between the short and long membrane-bound states and their relative populations. These resonances display only limited additional broadening of 3–4 Hz (approx. 10 *s*^−1^) compared to the vesicle-free spectrum (Figure 5C). This may be caused by life-time broadening due to exchange between the short and long membrane-bound states. To estimate the timescale of this exchange process and the relative populations of the different membrane-bound states, we simulated a three-state exchange system comprising the free state and two membrane-bound states spanning residues 1-16 (B16) or 1-100 (B100) (SI and Figure S9). The parameters extracted from the simulations are summarized in Figure 6A. Line-broadening similar to what we observed experimentally can be obtained with the following parameters: population of B16 between 5 and 20%, population of B100 54-69% (accordingly) and 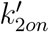 from B16 to B100 of 30-70 *s*^−1^. 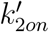 is a pseudo-first order association rate constant expressed as 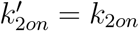. [binding sites]. Assuming 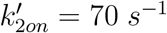 and that the population of the B16 state is 5%, the upper bound for the exchange rate constant between B16 and B100 expressed as 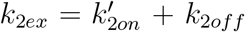 is 75 s^−1^ (see Supplementary Information). The lower bound for *k*_2*ex*_ is estimated for 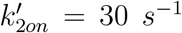 and the population of the B16 state of 20% and equals *k*_2*ex*_ = 41 *s*^−1^. Thus, we can conclude that the exchange between the free protein and short membrane-bound state occurs on a much shorter timescale than the exchange between short and long membrane-bound states. In addition, our simulations indicate that the relative population of the short membrane-bound state (B16 and B50 at L/P 50 and 100, respectively) can be substantial, which is expected to contribute to modulation of the properties of the *α*-synuclein-covered membrane interface.

**Figure 6.**
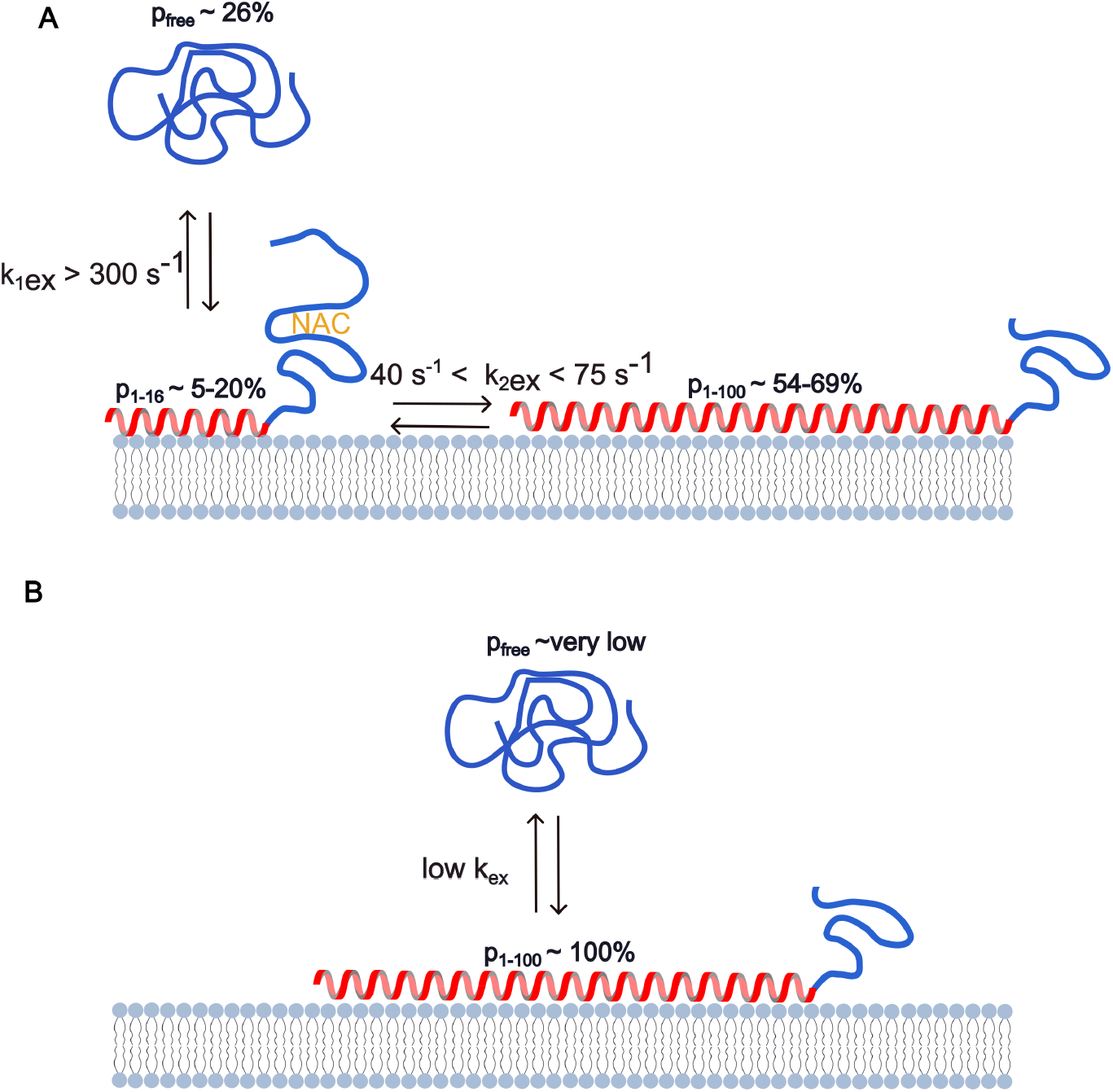
Cartoon summarizing the information about the presence of different protein states (free, short membrane-bound and long membrane-bound), their populations and timescales of exchange between them for L/P ratio 50, where *α*-synuclein amyloid formation is accelerated by lipid membranes (A) and L/P ratio 200 (B), where it is inhibited.

## Discussion

We carried out a detailed characterization of the equilibrium between free *α*-synuclein monomer and its membrane-bound state under conditions where lipid membranes either accelerate or inhibit amyloid formation. Interestingly, we find clear differences in the dynamics and heterogeneity of the protein-covered membrane interface in the two sets of conditions, which is likely to play an important role in modulating the free energy barrier for amyloid formation.

In conditions of protein excess (low L/P ratio), where *α*-synuclein amyloid formation is accelerated compared to the conditions without membranes, the protein-covered membrane is a dynamic and heterogeneous interface (Fig 6A). Apart from populating the free state, *α*-synuclein explores a range of membrane-bound states interacting with the membrane with sequence segments of different length (residues 1-16, 1-55 and 1-100). The free protein exchanges relatively fast (*k*_1*ex*_ ≥ 300 *s*^−1^) with the short membrane bound state and the short membrane-bound state exchanges on a longer timescale with the long membrane-bound state (*k*_2*ex*_ ≈ 75 *s*^−1^). The membrane is decorated with disordered segments of the adsorbed proteins of different length (segments 17-140, 56-140, 101-140). Thus, the highly hydrophobic region spanning residues 61-95 (NAC), which forms the core of *β*-sheets in *α*-synuclein amyloid fibrils is exposed to the surrounding solution and thus available to interact with other partly membrane-bound proteins or with the free monomer (Figure 6A).

In conditions of excess lipid membranes (high L/P ratio), where *α*-synuclein amyloid formation is inhibited, there is no detectable free protein in solution. In these conditions, virtually all protein populates the fully membrane-associated state, in which the NAC region is part of the long *α*-helix that is associated with the membrane (Fig 6B). Thus, at high L/P conditions, the protein-covered membrane interface is more homogeneous than at low L/P since all adsorbed proteins expose the disordered segments spanning residues 101-140 to the solution. We expect that the exchange process between the free and fully membrane-bound protein occurs on even longer timescale than the exchange between the short and long membrane-bound states (Figure 6B) based on an analogy to polymer desorption from a surface. The longer the adsorbed protein segment, the greater the number of protein-membrane contacts that need to be broken upon dissociation, which results in a low dissociation rate constant (*k*_*off*_). Also, strong positive cooperativity of *α*-synuclein association with lipid membranes may involve attractive protein-protein interactions.^26^ Such cohesive forces may constitute a kinetic barrier for desorption and thus decrease the *k*_*off*_.^27^

Altogether, the dynamics and heterogeneity of the *α*-synuclein-covered membrane interface is modulated by the L/P ratio. Modulation of the properties of the protein-covered membrane by the L/P ratio may explain the different effects that membranes have on *α*-synuclein amyloid formation in different L/P regimes, with the free energy barrier for amyloid formation being decreased in the low L/P ratio regime. It was previously inferred that interaction of an incoming monomer with the C-termini of membrane-associated proteins is critical in the acceleration of *α*-synuclein amyloid formation by lipid membranes.^28^ MD simulations revealed a preferred alignment of the monomer with respect to C-termini grafted on a surface, which is due to anisotropic charge distribution in the protein sequence and can be screened by increasing ionic strength.^28^ This electrostatic interaction is expected to be modulated by the properties of the protein segments extending from the membrane and thus by the L/P ratio. Moreover, the dynamics of the exchange between free and membrane-bound protein is also expected to play a role. Fast exchange at low L/P is likely to increase the frequency of encounters of the free monomer with the disordered segments of the bound proteins, which may lead to a nucleation event.

Inhibition of amyloid formation at high L/P ratio may be the result of either the fully membrane-bound state being the most thermodynamically stable state overall, considering also the fibrillar state (as was shown to be the case for *α*-synuclein-DMPS system^5^) or being a metastable state due to a high energy barrier separating it from amyloid fibrils. Regardless of which of the two is the case, the fully membrane-bound state of *α*-synuclein can be thought of having a protective role against aggregation.

The effect of membranes on *α*-synuclein amyloid formation can also be rationalized in terms of protein-membrane affinity. The interaction of the partly membrane-bound states with the membrane is weaker than the interaction of the fully membrane-bound state. Comparison of the HSQC data at 20 and 37^°^C (Figure 5B and Figure S4) shows that the presence of the short binding modes at a constant L/P ratio can be suppressed/induced by increasing/decreasing the overall strength of the protein-membrane interaction. The affinity of *α*-synuclein-membrane interaction can be modulated on the protein level (e.g., point mutations,^18,29,30^ post-translational modifications^31–33^ such as N-terminal acetylation), by changes in membrane properties (lipid composition in terms of length and saturation of acyl chains as well as headgroup chemistry),^8,30,34–36^ as well as by varying solution conditions (ionic strength, pH^16^). Changes in any of the above-mentioned parameters may trigger a shift from non-aggregating to aggregating conditions and thus from physiological to pathological *α*-synuclein-lipid self-assembly, or vice versa at a fixed L/P ratio.

The presence of different membrane-bound states under aggregating and non-aggregating conditions may be considered a manifestation of the Sabatier’s principle, which states that a catalyst (here, lipid membrane) does not work efficiently when the substrate (here, *α*-synuclein) binds too strongly. Indeed, a study of protein aggregation in the presence of different interfaces revealed that the heterogeneous primary nucleation rate follows a non-monotonic dependence on the protein-surface interaction strength, reaching a maximum for intermediate affinities.^37^ Too weak interaction results in insufficient concentration of protein on the interface, while too strong interaction is expected to interfere with the formation of growth-competent nuclei.

Altogether, our results emphasize the strong coupling of the equilibrium between free and membrane-bound *α*-synuclein and amyloid formation, which suggest a coupling between the physiological function of the protein and its aberrant aggregation. This coupling should be considered when designing therapeutic approaches for treating *α*-synuclein-related neurodegenerative diseases.

## Materials and Methods

### *α*-synuclein expression and purification

*α*-synuclein of human wild-type sequence, isotope-labelled with ^15^N or with a N122C mutation, were expressed in *E. coli* from Pet3a plasmids with *E. coli*-optimized codons (purchased from Genscript, Piscataway, New Jersey). The wild-type and ^15^N-labelled protein was purified using heat treatment, ion-exchange and gel filtration chromatography, as previously described.^38^ The N122C mutant was purified using the same protocol but with 1 mM dithiothreitol (DTT) included in all buffers. All experiments started with gel filtration on a 10×300 mm Superdex 75 column (GE Healthcare) to isolate fresh monomer in 10 mM MES buffer at pH 5.5.

### *α*-synuclein labelling

*α*-synuclein N122C mutant was labelled with Alexa Fluor 488 Malemide. Gel filtration was used to remove DTT from the protein and to exchange the buffer to 20 mM sodium phosphate, pH 8, supplemented with one molar equivalent of Alexa Fluor 488 Malemide dye from a 5 mM stock in DMSO, and incubated for 1 h. Excess free dye and phosphate buffer were removed using gel filtration on a 10×300 mm Superdex 75 column in 10 mM MES pH 5.5. In the text, the Alexa Fluor 488 labelled protein is referred to as *α*-synuclein-488.

### Lipids

The following liophilized lipids were purchased from Avanti Polar Lipids (Alabaster AL): 1,2-dioleoyl-sn-glycero-3-phospho-L-serine sodium salt (DOPS), 1,2-dioleoyl-sn-glycero-3-phosphocholine (DOPC). Liophilized Atto-647 1,2-Dipalmitoyl-sn-Glycero-3-

Phosphoethanolamine (DPPE-647) was purchased from ATTO-TEC GmbH.

### SUV preparations

Small unilamellar vesicles (SUVs) were prepared by extrusion using Avanti Mini Extruder (Avanti Polar Lipids). The desired volume of 7:3 (molar ratio) DOPC:DOPS mixture in chloroform:methanol (7:3 volume ratio) was left overnight in a vacuum oven at room temperature for the solvent to evaporate. The dried lipids were then hydrated with 10 mM MES buffer at pH 5.5 and left on stirring for 2 h at room temperature. The 100 nm pore size filters were first saturated with lipid solution that was discarded. The SUVs were then obtained by extruding 21 times.

### Aggregation kinetics

20 *µ*M *α*-synuclein was incubated in 10 mM MES pH 5.5, 0.01% *NaN*_3_ in presence of 20 *µ*M ThT and increasing concentrations of SUVs in a 96-well non-binding plate (PEGylated polystyrene, Corning, 3881). The ThT fluorescence signal was monitored as a function of time using a plate reader (FluoStar Omega, BMG Labtech, Offenburg, Germany) under quiescent conditions at 37 ^°^C with excitation at 440 nm and emission at 480 nm.

### Agarose gel electrophoresis

Agarose gel electrophoresis was run in horisontal mode using a home-built apparatus as described.^39^ All the samples and the gels (1% (w/v) Sea Kem LE Agarose from LONZA Bioscience) were prepared in 50 mM MES pH 5.5. All samples contained 20 *µ*M *α*-synuclein-488 and varying amounts of DOPC:DOPS 7:3 SUVs with 2% of Atto-647-DHPE. 15 *µ*l of each sample was placed in the agarose gel wells. A potential of 240 V was applied and the samples were allowed to migrate in the gel for 20 minutes. SUVs used for the agarose gel electrophoresis experiment were extruded through 50 nm filters. The size distribution and polydispersity index were analyzed using Malvern Zetasizer Nano-Z (Malvern Instruments Ltd.). The average hydrodynamic radius of SUVs was 37 nm and polydispersity index 0.06.

### Circular Dichroism Spectroscopy

The samples were prepared by incubating *α*-synuclein with DOPC:DOPS 7:3 SUVs in 10 mM MES buffer, pH 5.5, 0.01% *NaN*_3_. *α*-synuclein concentration was in the range 2-10 *µ*M and lipid concentration was varied accordingly. The overall concentration was adjusted in order to obtain good quality spectra at all analyzed L/P ratios. Far-UV spectra were recorded using a JASCO J-815 equipped with a Peltier type cell holder using 2 mm quartz cuvettes. Each CD spectrum was an average over 5 individual spectra recorded between 250 nm - 200 nm, with a bandwith of 1 nm, a data pitch of 1 nm, a scanning speed of 10 nm/min and an integration time of 8 s. For each sample the CD signal of the buffer was subtracted from the CD signal of the protein.

### PFG NMR

PFG NMR spectroscopy experiments were performed at 25^°^C using a Bruker Avance Neo 500 MHz NMR spectrometer equipped with a MIC gradient probe with a 10 mm ^1^H insert. The pulse sequence used was diffSte from the Bruker library. The strengths of 5.2 ms long gradient pulses were incremented logarithmically in 64 steps to the maximal value of 136, 150, 180, 295 and 280 Gauss/cm for samples at L/P 0, L/P 50, L/P 100, L/P 200 and free SUVs, respectively. For all measurements, the diffusion time was kept constant at 50 ms and a short pre-acquisition delay of 10 *µ*s was used.

The SUVs for PFG STE experiments were prepared in MES/NaOD buffer in D_2_O. The pH of the MES buffer was adjusted with NaOD to the value of 5.5 (reading of the pH meter). Wild-type *α*-synuclein was isolated by gel filtration on a 10×300 mm Superdex 75 column in 10 mM MES/NaOH pH 5.5, desalted using a 16×25 mm HiTrap Desalting column and lyophilized. Immediately prior to the experiment, the protein was resuspended in the SUV dispersion in MES/NaOD to achieve a protein concentration of 200 *µ*M.

### ^1^H-^15^N Heteronuclear Single Quantum Coherence experiment

The HSQC experiments were carried out for 20 *µ*M ^15^N-labelled *α*-synuclein alone and with SUVs at different L/Ps. The experiments at 20 and 37 °C were acquired using a Varian VNMR-S 600 MHz and 500 MHz spectrometer, respectively, equipped with room temperature inverse probes, and the pulse sequence gNhsqc. All spectra were recorded with spectral widths of 10 ppm and 28 ppm in the ^1^H and ^15^N dimension, respectively, covered by 1458 and 128 points, using a recycle delay of 1 s and an acqusition time of 121 ms. Typically 48 scans were acquired. All spectra were processed using NMRpipe^40^ employing a processing protocol with solvent filter, square cosine apodization and zero filling to double the number of points in all dimensions. Peak volumes, line widths and intensities were evaluated using the program suite PINT.^22,23^ For analysis of the HSQC spectra, we transferred published peak assignments by comparing peak positions from Bodner et al^29^ and Vandova et al., BMRB Entry 18857.

## Acknowledgements

This work was supported by the Knut and Alice Wallenberg Foundation (2016.0074 to SL and ES) and by the Swedish Research Council (2019-02397 to SL and ES).

## Supporting Information Available

Additional ^1^*H*-^15^*N* Heteronuclear Single Quantum Coherence spectroscopy and Circular Dichroism spectroscopy data, description of the analysis of PFG NMR data and simulations of NMR spectra due to exchanging states.

