## Supporting Information for "Tipping point in *α*-synuclein–membrane interactions: stable protein-covered vesicles or amyloid aggregation"

‡*Centre for Analysis and Synthesis, Center for Chemistry and Chemical Engineering, Lund University, SE-22100 Lund, Sweden*

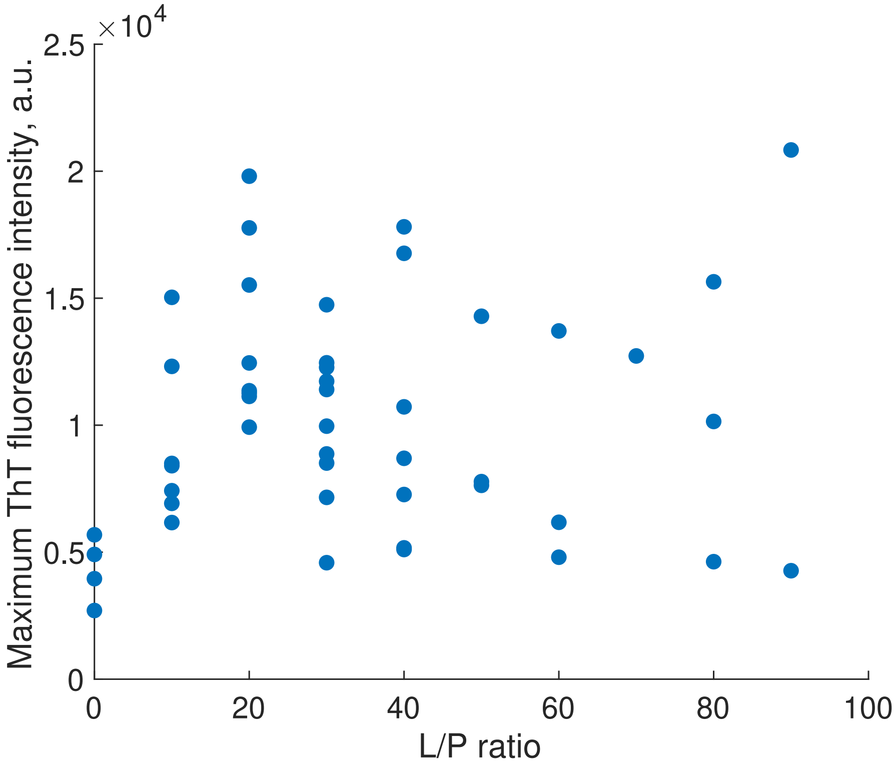


Figure S1: Maximum ThT fluorescence intensity in the aggregation kinetics assay shown in Figure 1 as a function of L/P ratio. The $\alpha$-synuclein concentration was kept constant at 20 $\mu$M, while the lipid concentration was varied. No ThT signal above the baseline was detected for L/P $>$ 100.

### Calculation of free and bound protein fractions from diffusion coefficients extracted from PFG-diffusion NMR

The diffusion coefficient of $\alpha$-synuclein in samples containing $\alpha$-synuclein and vesicles, $D_{average}$, is a population average of the diffusion coefficients of the free and vesicle-bound protein. From the diffusion coefficients presented in Table 1, we calculated the fraction of vesicle-bound $\alpha$-synuclein at different L/P ratios, $x$, according to the equation:

$$x = \frac{D_{average} - D_{free}}{D_{freeSUV} - D_{free}} \tag{1}$$

$D_{average}$ is the diffusion coefficient extracted from the decay of the signal at 7.2 ppm in the samples containing protein and vesicles, $D_{free}$ is the diffusion coefficient extracted from the decay of the signal at 7.2 ppm in the free protein sample and $D_{vesicle}$ is the diffusion coefficient extracted from the decay of the signal at 1 ppm in the free SUV sample (Figure 4).

Table S1: Diffusion coefficients extracted from the intensity decay curves of the peaks corresponding to free $\alpha$-synuclein and $\alpha$-synuclein in the presence of SUVs (7.2 ppm), free or protein-bound SUVs (1 ppm) and MES molecule (3.5 ppm). In the case of samples containing both $\alpha$-synuclein and SUVs, the diffusion coefficient extracted from the signal at 7.2 ppm corresponds to a population average of the free and membrane-bound protein states.

| Sample | Peak | $D_{coeff}$ $m^2/s$ |
|---|---|---|
| free $\alpha$-synuclein | 7.2 ppm | $7.3 \cdot 10^{-11}$ |
| SUVs + $\alpha$-synuclein at L/P 50 | 7.2 ppm | $2.4 \cdot 10^{-11}$ |
| SUVs + $\alpha$-synuclein at L/P 100 | 7.2 ppm | $1.0 \cdot 10^{-11}$ |
| SUVs + $\alpha$-synuclein at L/P 200 | 7.2 ppm | $3.0 \cdot 10^{-12}$ |
| free SUVs | 1 ppm | $6.6 \cdot 10^{-12}$ |
| MES | 3.5 ppm | $6.0 \cdot 10^{-10}$ |

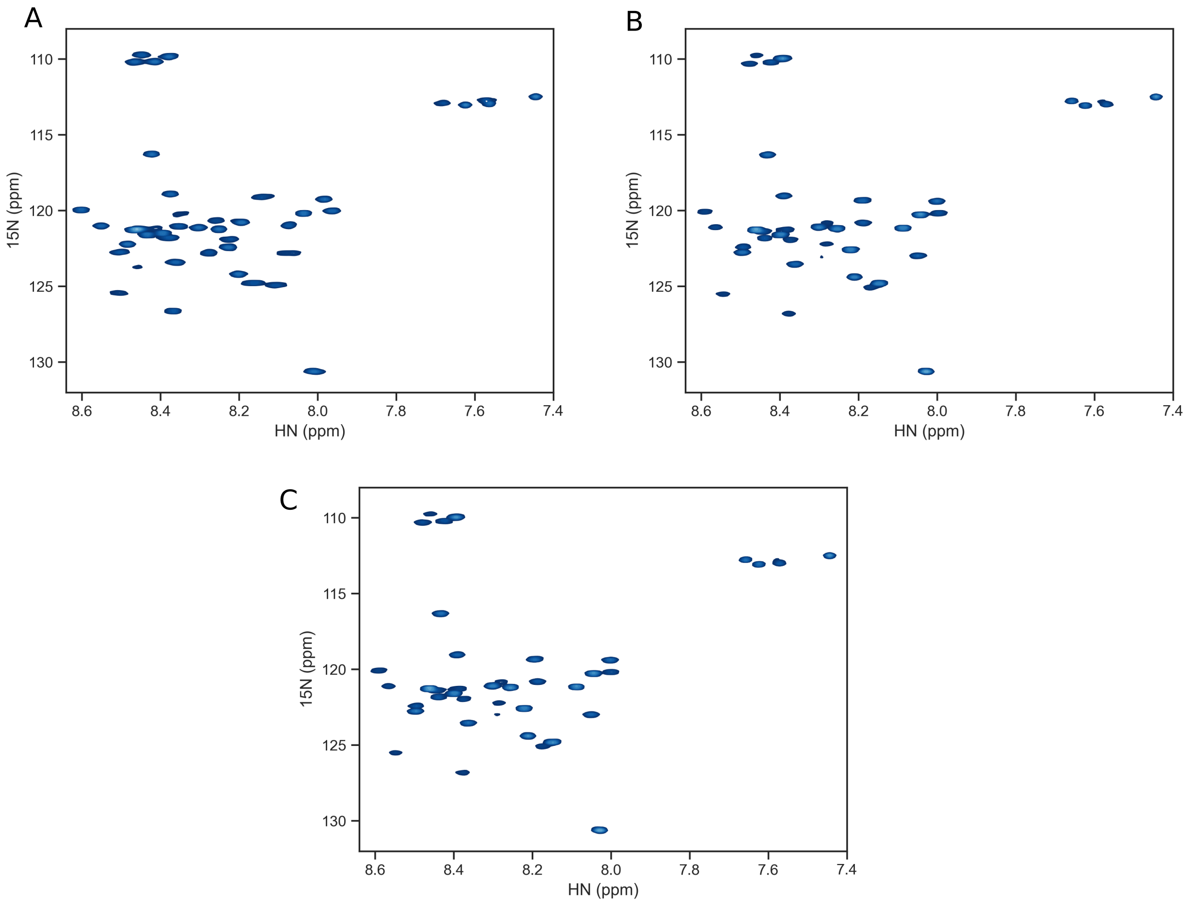


Figure S2: $^{1}$H-$^{15}$N HSQC spectra at 20°C. A) $\alpha$-synuclein in the presence of DOPC:DOPS 7:3 SUVs at L/P 100. B) $\alpha$-synuclein in the presence of DOPC:DOPS 7:3 SUVs at L/P 200. C) $\alpha$-synuclein in the presence of DOPC:DOPS 7:3 SUVs at L/P 250.

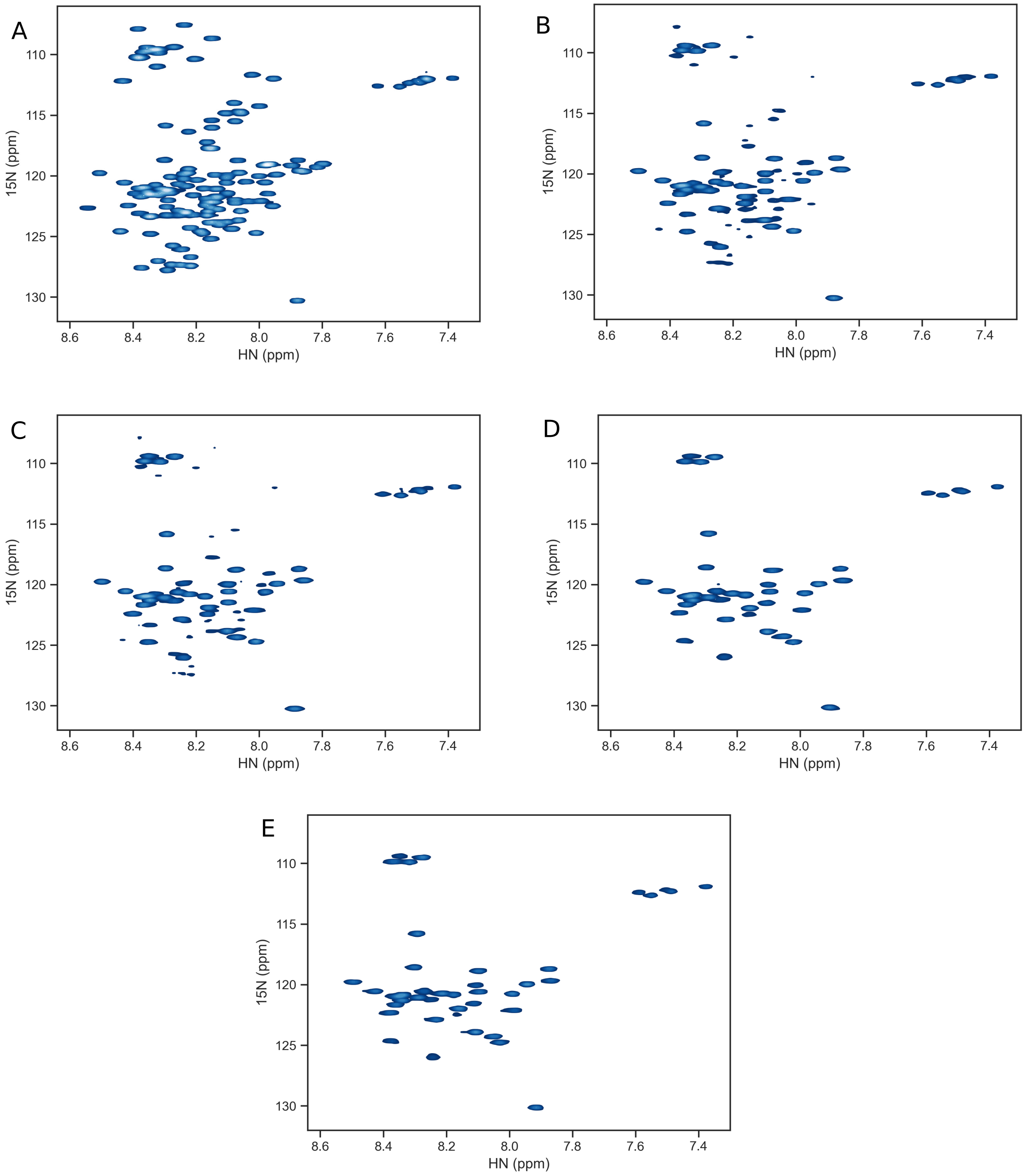


Figure S3: $^{1}H$-$^{15}N$ HSQC spectra at 37°C. A) Free $\alpha$-synuclein. B) $\alpha$-synuclein in the presence of DOPC:DOPS 7:3 SUVs at L/P 10. C) $\alpha$-synuclein in the presence of DOPC:DOPS 7:3 SUVs at L/P 20. D) $\alpha$-synuclein in the presence of DOPC:DOPS 7:3 SUVs at L/P 40. E) $\alpha$-synuclein in the presence of DOPC:DOPS 7:3 SUVs at L/P 50.

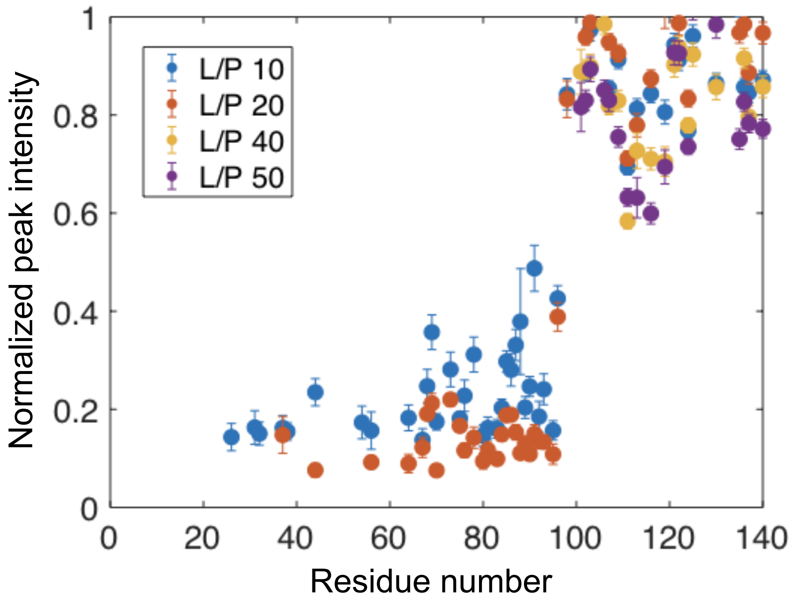


Figure S4: $\alpha$-synuclein binding to DOPC:DOPS 7:3 SUVs studied with $^{1}H$-$^{15}N$ HSQC NMR at 37°C. Peak intensities in the samples containing $\alpha$-synuclein and SUVs at L/P 10, 20, 40 and 50, normalized with respect to the intensity of the corresponding peaks in the free $\alpha$-synuclein spectrum and plotted as a function of the residue number in the $\alpha$-synuclein sequence.

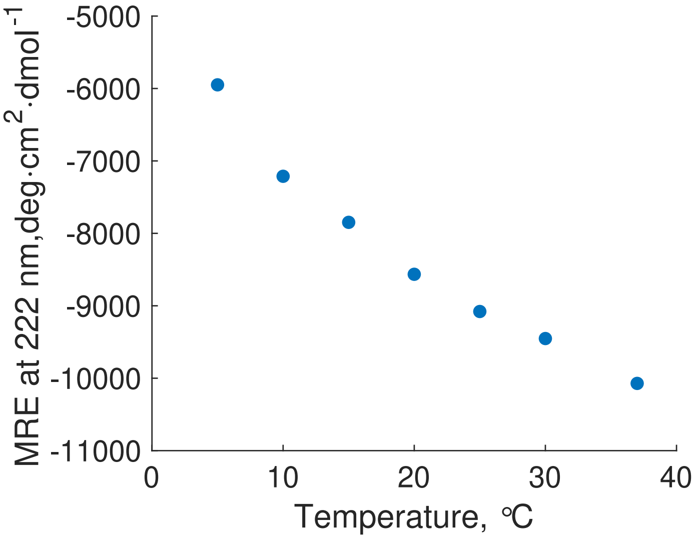


Figure S5: Temperature dependence of $\alpha$-synuclein association with lipid membranes as studied using circular dichroism spectroscopy. Mean residue ellipticity at 222 nm as a function of temperature in the 5-37°C temperature range for a sample containing $\alpha$-synuclein and DOPC:DOPS 7:3 SUVs at L/P 50.

### Simulation of Exchange Spectra

The NMR spectrum of an exchanging system can be simulated by solving the Bloch-McConnell equations,[1] see e.g. ref.[2] or ref.[3]

#### Effect of $\Delta R_2$ for two state exchange

Consider an exchange between two states:

$$\mathrm{F} \xrightleftharpoons{k_{ex}} \mathrm{B16}$$

where F represents the free protein state, B16 the state where amino acid residues 1-16 are bound to the vesicle, and $k_{ex} = k'_{on} + k_{off}$ is the exchange rate constant. For binding of $\alpha$-synuclein to vesicles, $k'_{on}$ should be interpreted as a pseudo-first order rate constant, $k'_{on} = k_{on}C_{site}$, where $C_{site}$ is the concentration of available binding sites, which will be different for the different L/P ratios. Let $p_F$ and $p_B$ represent the fractions of free and bound protein, respectively, and $\nu_F$ and $\nu_B$ be the resonance frequencies for the free and bound states. Then we have

$p_B = 1 - p_F$,

$K'_a = p_B/p_F = k'_{on}/k_{off}$,

$k'_{on} = K'_a k_{ex}/(1 + K'_a)$, and

$k_{off} = k_{ex}/(1 + K'_a)$.

The kinetic matrix $\mathbf{K} = \begin{pmatrix} -k'_{on} & k_{off} \\ k'_{on} & -k_{off} \end{pmatrix}$,

the relaxation matrix $\mathbf{R_2} = \begin{pmatrix} R_{2F} & 0 \\ 0 & R_{2B} \end{pmatrix}$,

where $R_{2F}$ and $R_{2B}$ are the transverse relaxation rate constants for state F and B16, and the chemical shift matrix $\omega_{\mathbf{0}} = \begin{pmatrix} \omega_{0F} & 0 \\ 0 & \omega_{0B} \end{pmatrix}$,

where $\omega_{0F} = 2\pi\nu_F$ and $\omega_{0B} = 2\pi\nu_B$. From the combined matrix $\mathbf{L} = i \cdot \omega_{\mathbf{0}} - \mathbf{R_2} + \mathbf{K}$, the free induction decay can then be calculated, and Fourier transformed to obtain the spectrum.[3] Figure S6A shows simulated spectra for symmetric exchange ($k'_{on} = k_{off}; p_F = p_B$) with different values of $k_{ex}$ and $R_{2F} = R_{2B}$. There is a clear progression from slow to fast exchange, relative to the chemical shift difference, as $k_{ex}$ increases.

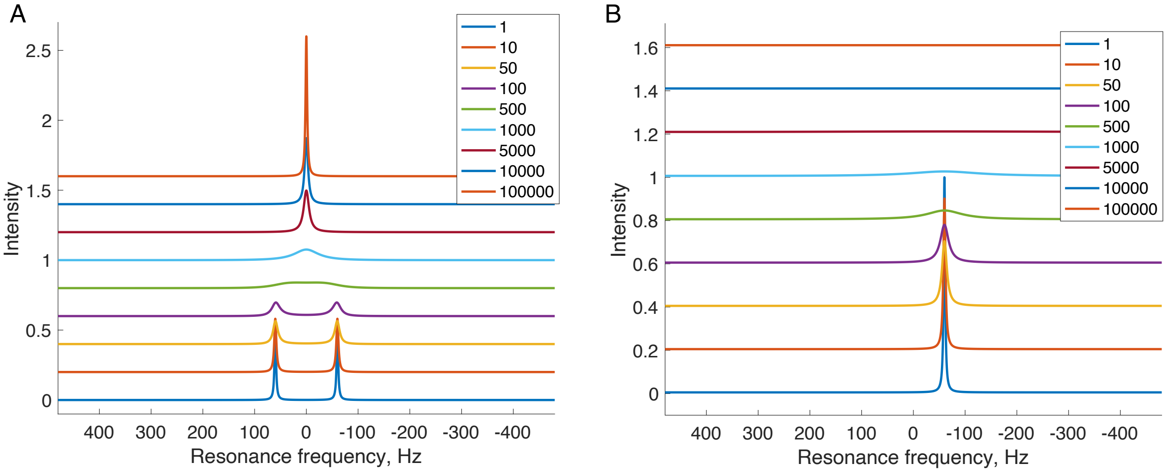


Figure S6: Simulated spectra for symmetric exchange between two spin states with different values of $k_{ex}$. The resonance frequencies of the free and bound state are set to $\nu_F$ = -60 Hz, $\nu_B$ = 60 Hz. The different values used for $k_{ex}$ are given in the insert in units of $s^{-1}$. In panel A, the transverse relaxation rate constants for both states are the same, $R_{2F} = R_{2B} = 10$ $s^{-1}$, while in panel B, $R_{2F} = 10$ $s^{-1}$, $R_{2B} = 39\ 000$ $s^{-1}$.

The NMR spectra of a system with a large difference in the transverse relaxation rates of the two sites will be much different from the case described above. Figure S6B shows simulated spectra for a system with the same parameters as in Figure S6A, except that $R_{2B} = 39\ 000$ $s^{-1}$. In contrast to the exchange spectra obtained when $R_2$ are of similar values for the two sites, it is clear that the chemical shift of the observable resonance does not change, but will be broadened beyond detection at large values of $k_{ex}$ due to life-time broadening. These facts are not always considered when analyzing NMR data of an exchanging system, *e.g.* where a small molecule associates with a much larger entity, and can lead to erroneously determined limits for the exchange rate if the difference in chemical shifts are used to define the NMR time-scale. As an example, in Figure S7, we show simulated spectra for a system close to coalescence ($k_{ex} \approx \Delta\omega_0$), with increasing values of $R_{2B}$. Again, at large values of $R_{2B}$, the chemical shift of the observed resonance will appear at the frequency of the free state, which could be misinterpreted as $k_{ex} \ll \Delta\omega_0$ if the large difference in $R_2$ is not taken into account.

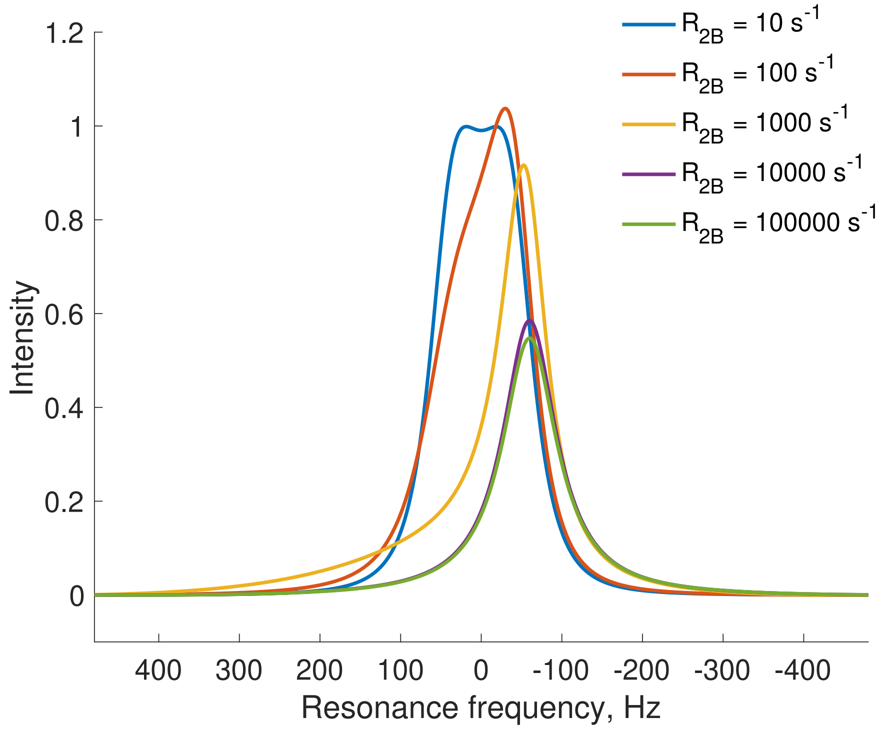


Figure S7: Simulated spectra for symmetric exchange for different values of the transverse relaxation rate of the bound state, $R_{2B}$. The resonance frequencies of the free and bound state are set to $\nu_F$ = -60 Hz, $\nu_B$ = 60 Hz, and $k_{ex}$ = 500 $s^{-1}$. The different values used for $R_{2B}$ are given in the insert in units of $s^{-1}$.

#### Estimating a lower bound for $k_{ex}$

Simulated NMR spectra can provide an estimate of the lower bound of the exchange rate constant for the binding of free protein to the vesicles. We know from the PFG NMR data that the fraction of free protein is $p_F \sim 26\%$ at L/P 50. Still, the $^{15}$N-HSQC peaks from the N-terminal part of the protein are absent, which indicate that they are broadened beyond detection. Figure S5A shows simulated spectra for this case: asymmetric exchange with $p_B/p_F = 3$, $R_{2F} = 10\ \mathrm{s}^{-1}$, and $R_{2B} = 39\ 000\ \mathrm{s}^{-1}$, for different values of $k_{ex}$. For exchange rates $k_{ex} \geq 300\ \mathrm{s}^{-1}$ the resonance from the free state will be sufficiently life-time broadened to be undetectable. However, the lower limit for $k_{ex}$ depends on the relative populations of the two states, so that a lower value of $p_B/p_F$ will further increase the lower limit of $k_{ex}$, as exemplified in Figure S5B. Note that the results are insensitive to the value of $\Delta\omega_0$ as long it is much smaller than $\Delta R_2$.

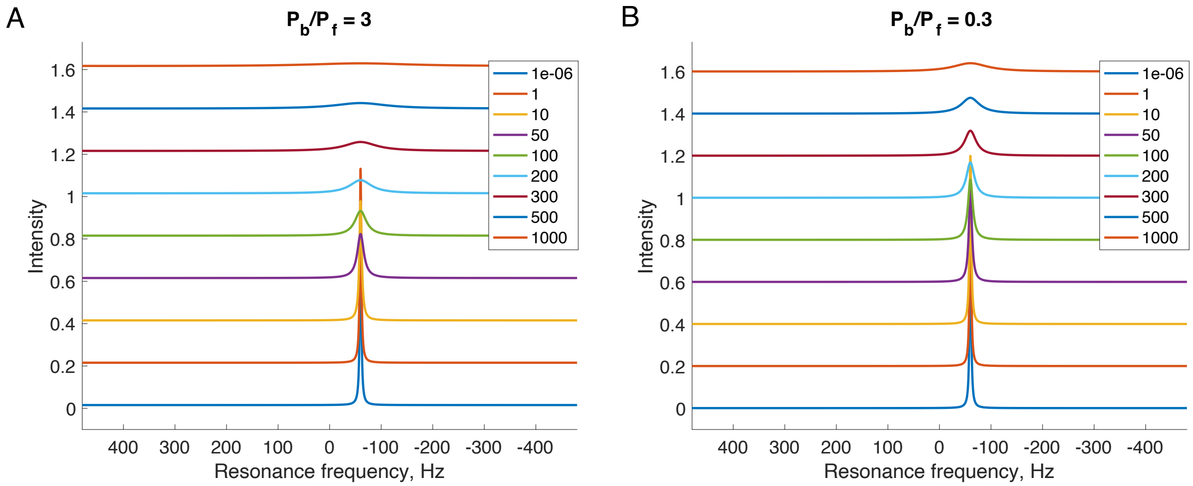


Figure S8: Simulated spectra for asymmetric exchange with different values of $k_{ex}$. The resonance frequencies of the free and bound state are set to $\nu_F$ = -60 Hz, $\nu_B$ = 60 Hz and the transverse relaxation rates of the free and bound state are $R_{2F} = 10\ \mathrm{s}^{-1}$, and $R_{2B}$ = 39 000 $\mathrm{s}^{-1}$. In panel A, the population ration of the bound and free states, $p_B/p_F = 3$, while in panel B, $p_B/p_F = 1/3$. The different values used for $k_{ex}$ are given in the inserts in units of $\mathrm{s}^{-1}$.

### Three State Exchange

For L/P 50 sample we consider the following exchange between three states:

$$\mathrm{F} \underset{}{\overset{k_{1ex}}{\rightleftharpoons}} \mathrm{B16} \underset{}{\overset{k_{2ex}}{\rightleftharpoons}} \mathrm{B100}$$

where F represents the free protein state, and B16 and B100 represent the states where amino acid residues 1-16 and 1-100, respectively, are associated with the vesicle. $k_{1ex} = k'_{1on} + k_{1off}$ and $k_{2ex} = k'_{2on} + k_{2off}$ are the exchange rate constants for exchange between F and B16, and between B16 and B100, respectively. Again, $k'_{1on}$ and $k'_{2on}$ are the pseudo-first order rate constants. The fractions of the three states are denoted $p_F$, $p_{16}$, and $p_{100}$. Then we have

$p_{100} = 1 - p_F - p_{16}$,

$K'_{1a} = p_{16}/p_F$,

$k'_{1on} = K'_{1a} k_{1ex}/(1 + K'_{1a})$,

$k_{1off} = k_{1ex}/(1 + K'_{1a})$, and

$k_{2off} = p_{16}/p_{100} \cdot k'_{2on}$.

The kinetic matrix $\mathbf{K} = \begin{pmatrix} -k'_{1on} & k_{1off} & 0 \\ k'_{1on} & -k_{1off} - k'_{2on} & k_{2off} \\ 0 & k'_{2on} & -k_{2off} \end{pmatrix}$.

Now assume that the chemical shift of an amino acid not in direct contact with the vesicle is $\omega_{0F}$, i.e. the chemical shifts of amino acids 17-100 in state B16 are the same as in the free state. When the amino acid is in direct contact with the vesicle, the chemical shift changes with $\Delta\omega$. Also, assume that the transverse relaxation rate of an amino acid not in direct contact with the vesicle is $R_{2F}$, i.e. $R_2$ of amino acids 17-100 in state B16 is the same as in the free state. When the amino acid is in direct contact with the vesicle, $R_2$ changes to $R_{2B}$. For amino acids 1-16, the chemical shift matrix $\omega_\mathbf{0}$ and the relaxation matrix $\mathbf{R_2}$ are given by

$$\omega_\mathbf{0} = \begin{pmatrix} \omega_{0F} & 0 & 0 \\ 0 & \omega_{0F} + \Delta\omega & 0 \\ 0 & 0 & \omega_{0F} + \Delta\omega \end{pmatrix},$$

$$\mathbf{R_2} = \begin{pmatrix} R_{2F} & 0 & 0 \\ 0 & R_{2B} & 0 \\ 0 & 0 & R_{2B} \end{pmatrix}.$$

For amino acids 17-100, the chemical shift matrix $\omega_\mathbf{0}$ and the relaxation matrix $\mathbf{R_2}$ are given by

$$\omega_{\mathbf{0}} = \begin{pmatrix} \omega_{0F} & 0 & 0 \\ 0 & \omega_{0F} & 0 \\ 0 & 0 & \omega_{0F} + \Delta\omega \end{pmatrix},$$

$$\mathbf{R_2} = \begin{pmatrix} R_{2F} & 0 & 0 \\ 0 & R_{2F} & 0 \\ 0 & 0 & R_{2B} \end{pmatrix}.$$

For amino acids 101-140, the chemical shift matrix $\omega_{\mathbf{0}}$ and the relaxation matrix $\mathbf{R_2}$ are given by

$$\omega_{\mathbf{0}} = \begin{pmatrix} \omega_{0F} & 0 & 0 \\ 0 & \omega_{0F} & 0 \\ 0 & 0 & \omega_{0F} \end{pmatrix},$$

$$\mathbf{R_2} = \begin{pmatrix} R_{2F} & 0 & 0 \\ 0 & R_{2F} & 0 \\ 0 & 0 & R_{2F} \end{pmatrix}.$$

We can allow for $\omega_{0F}$ to be different for amino acids 1-16, 17-100, and 101-140 so as to separate the resonances arising from the different protein regions. Setting $p_F = 0.26$ (from PFG NMR data) and $k_{1ex} = 300$ s$^{-1}$, we simulated NMR spectra to estimate values for $p_{16}$ and $k'_{2on}$. For each simulated spectrum in Figure S6, $p_{16}$ was chosen, and $k'_{2on}$ was varied until the spectrum matched the experimentally observed intensity of $\sim 17\%$ and line broadening ($R_{2,ex}$) of $\approx 10$ s$^{-1}$ were obtained for the resonances from amino acids 17-100. Good agreement with experimental data is reached for values of $p_{16} = 0.05$–$0.2$ and $k'_{2on} =$ 30–70 s$^{-1}$. Examples of parameters, which could not model our experimental data well are plotted in Figure S10. A too small value for $k'_{2on}$ results in a too small $R_{2,ex}$ contribution and too high intensity of the B100 peak, and a too large value for $k'_{2on}$ gives the opposite, larger $R_{2,ex}$ and smaller intensity of the B100 peak.

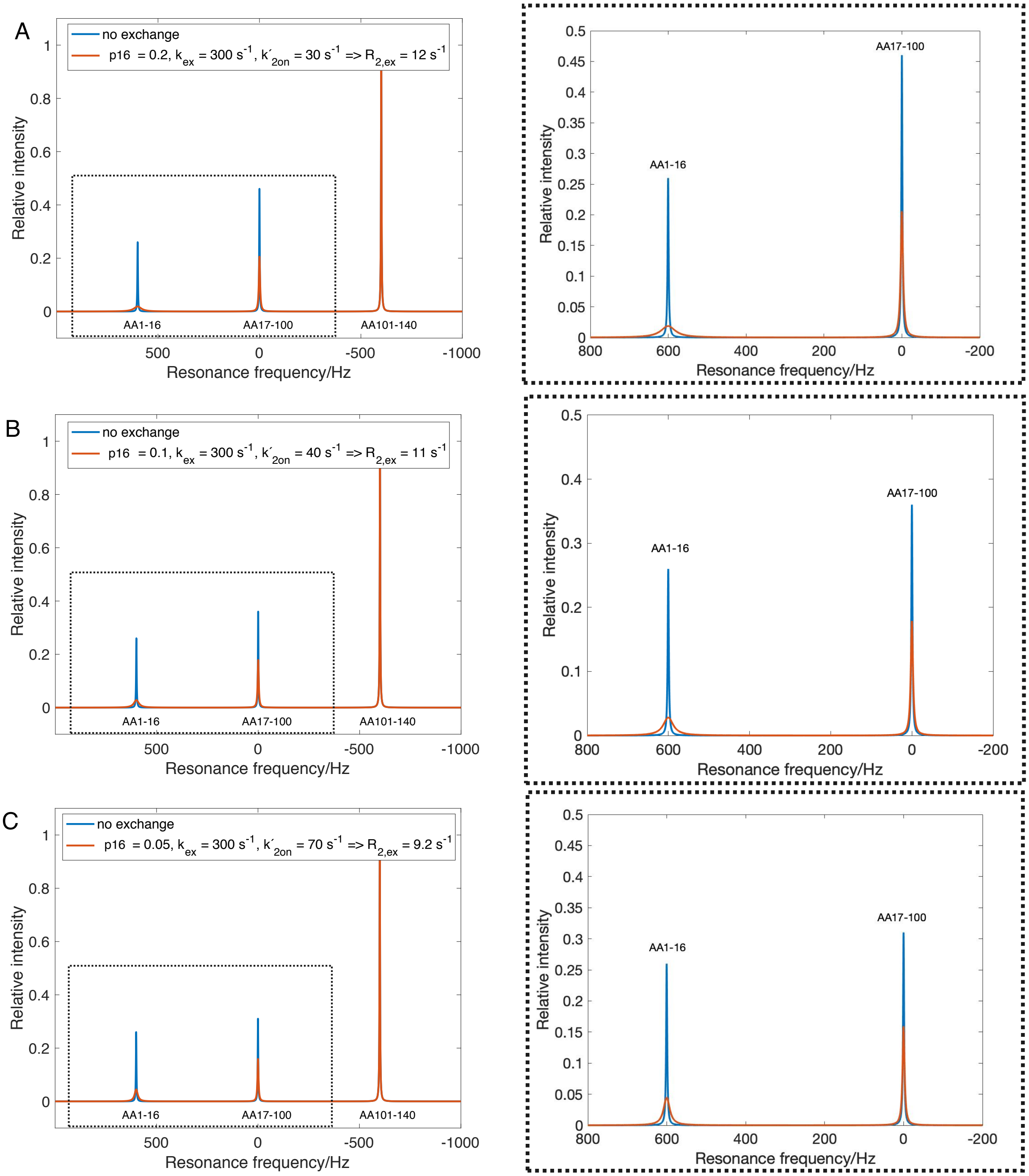


Figure S9: Three state exchange NMR spectra simulated to extract the values of the population of the short membrane-bound state, $p_{16}$, and the pseudo first-order association rate constant from the short to the long membrane-bound state, $k'_{2on}$, that can model the experimental data. The exchange rate constant between the free state and B16 was fixed to $k_{1ex} = 300$ s$^{-1}$, the free protein fraction to $p_F = 0.26$ and the transverse relaxation rates of the free and the membrane-bound states to $R_{2F} = 10$ s$^{-1}$, $R_{2B} = 39\,000$ s$^{-1}$. $\Delta\omega = 2\pi{\cdot}120$ s$^{-1}$. The obtained values for $p_{16}$ and $k'_{2on}$, are given in the inserts in the spectra. Thus, the experimentally measured broadening of the resonances corresponding to residues 17-100 of 10 $s^{-1}$ (Figure 5) corresponds to $p_{16}$ of 5-20% and $k'_{2on}$ 30-70 $s^{-1}$.

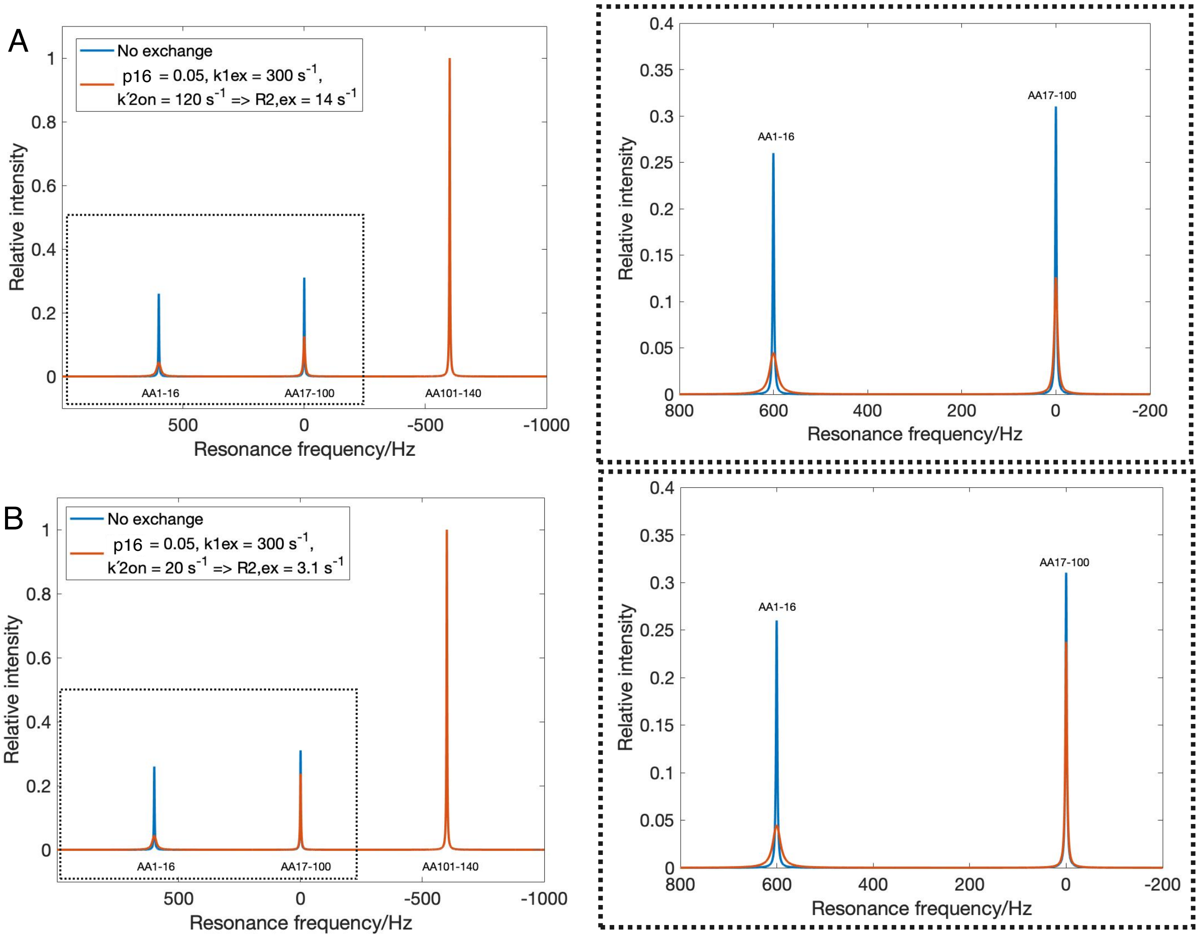


Figure S10: Examples of simulation parameters that could not model the experimental data. A too large value for $k'_{2on}$ results in a too large $R_{2,ex}$ contribution ($14\ s^{-1}$) and too low intensity of the B100 peak (A), while a too small value for $k'_{2on}$ results in a too small $R_{2,ex}$ contribution and too high intensity of the B100 peak (B).

(3) Waudby, C. A.; Alfonso, I. An introduction to one-and two-dimensional lineshape anal-

ysis of chemically exchanging systems. *Journal of Magnetic Resonance Open* **2023**, *16*, 100102.
